# Prevalence of Slam-dependent hemophores in Gram-negative bacteria

**DOI:** 10.1101/2023.03.29.534853

**Authors:** Hyejin Esther Shin, Chuxi Pan, David M. Curran, Thomas J. Bateman, Derrick HY Chong, Dixon Ng, Megha Shah, Trevor F. Moraes

**Affiliations:** Department of Biochemistry, University of Toronto, Toronto, Ontario, Canada

## Abstract

Iron acquisition systems are crucial for pathogen growth and survival in iron-limiting host environments. To overcome nutritional immunity, bacterial pathogens evolved to use diverse mechanisms to acquire iron. Here, we examined a heme acquisition system driven by hemophores called HphAs from several Gram-negative bacteria. Structural determination of HphAs revealed a N-terminal clamp-like domain that binds heme and a C-terminal eight-stranded β-barrel domain that shares the same architecture as the Slam-dependent Neisserial surface lipoproteins. The structure of these HphAs is strikingly similar to a novel hemophore discovered by Latham et al. (2019), named hemophilin^1^. The genetic organization of HphAs consist of genes encoding a Slam homolog and a TonB-dependent receptor (TBDR). We investigated the Slam-HphA system in the native organism or the reconstituted system in *E. coli* cells and found that the efficient secretion of HphA is dependent on Slam. The TBDR also played an important role for heme uptake and conferred specificity for its cognate HphA. Furthermore, bioinformatic analysis of HphA homologs revealed that HphAs are conserved in the alpha, beta, and gammaproteobacteria Together, these results show that HphA presents a new class of hemophores in Gram-negative bacteria and further expands the role of Slams in transporting soluble proteins supporting it role as a type 11 secretion system.

## Introduction

Iron is an essential nutrient that is important for many fundamental processes such as DNA synthesis and cellular respiration for nearly all living organisms. However, high levels of free iron can generate toxic reactive oxygen species (ROS) through the Fenton reaction and cause oxidative damage to macromolecules such as DNA, proteins and lipids^2^. To maintain a strict iron homeostasis, most of the free iron is incorporated into heme and bound to hemoproteins such as hemoglobin, myoglobin and cytochromes^3^. The remaining free iron is further stored into proteins including transferrin, lactoferrin, and ferritin. Restricting iron through these iron-binding proteins keeps the free iron concentration in the serum of about 10^−24^M which severely limits the amount of bioavailable iron for invading pathogens, a defense strategy known as nutritional immunity^4, 5^.

To survive in such iron-restricted environments, Gram-negative bacteria have evolved elaborate methods to hijack iron from the host. The most prevalent iron acquisition mechanism in bacteria is the production of siderophores. Siderophores are secondary metabolites produced by bacteria that bind ferric iron with high affinity (Kd, between 0.1 to 100 nM)^6^. The secretion of siderophores is facilitated by at least one or more efflux pump superfamilies depending on the siderophore type^7–9^. In order to retrieve iron from these siderophores, bacteria employ specific outer membrane proteins (OMPs) to bring the iron loaded siderophores back into the cell. In addition, bacteria utilize homologous outer membrane (OM) receptors to acquire iron from sources such as transferrin, lactoferrin, ferritin, heme, and/or hemoproteins. Iron acquisition through OM receptors often involves a TonB-dependent receptor (TBDR) and/or surface lipoproteins (SLPs). TBDRs consist of 22-stranded β-barrel domain and N-terminal plug domain. Extracellular loops of TBDR facilitate the binding with the substrate^10^. TBDRs transport nutrients into the bacterial cell using the energy coupled to the proton motive force harnessed across the inner membrane by the TonB-ExbB-ExbD complex^11^. Surface lipoproteins are a group of peripheral membrane proteins with unique functions including nutrient acquisition, immune evasion, and cell adhesion. A subset of SLPs, such as transferrin binding protein B (TbpB), are part of the bipartite receptor along with TBDRs to effectively uptake nutrients from the cell surface. Surface display of these lipoproteins are mediated by the surface lipoprotein assembly modulator (Slam)^12^.

Another strategy, restricted to heme iron sources, is to scavenge exogenous heme in the host environment by secreted proteins termed hemophores. However, approximately 67% of heme iron is bound to hemoglobin in the human body which severely limits the amount of intracellular free heme levels at roughly 0.1 µM for invading pathogens^13^. Several hemophores in Gram-negative bacteria have been reported, each with a unique fold and structural elements comprising the heme-binding site^1, 14–17^. Some of these hemophores were reportedly secreted by the Type 1 Secretion System (T1SS) and two-partner secretion system (TPS; also known as the Type Vb secretion system)^15, 18^.

In a recent report, a Slam homolog called HsmA was also shown to secrete a hemophore called hemophilin (HphA) in *Acinetobacter baumannii*^19^. Previous findings have shown that Slams are prevalent in Gram-negative bacteria but had only shown to be associated with the surface display of lipidated proteins, SLPs, until now^20^.Inspired by this finding, in this study, we performed bioinformatics on Slam-dependent hemophilins homologs and identified 1550 unique protein accessions mainly from the gamma-proteobacteria, and some in the alpha-and beta-proteobacteria. We report three additional structures of HphAs, belonging to the gammaproteobacteria, in *Stenotrophomonas maltophilia*, *Vibrio harveyi*, and *Haemophilus parainfluenzae*.

The structural characterization of the HphA structures from these organisms revealed the individual heme coordination mechanism and a common C-terminal eight-stranded β-barrel domain shared with other Slam-dependent surface lipoproteins^21^. We confirmed that the secretion of these HphAs is dependent on the expression of Slam. Finally, we investigated the importance and specificity of HphAs as an iron source using *in vivo* growth assays in *A. baumannii*.

## Results

### HphAs genes are located adjacent to Slam and TBDR genes

While searching for novel SLP substrates translocated by Slam we discovered a larger number of HphA-type hemophores. Previous analysis of gene neighbourhoods around the Slam homologs indicated the presence of genes predicted to encode lipoproteins and TBDRs^20^. However, we noticed that a subset of Slam genes were adjacent to genes that were predicted to be non-lipidated proteins. This observation was in agreement with the bioinformatic analysis performed by Grossman et al. (2021) where a cluster of Slams are found adjacent to proteins predicted to be processed by signal peptidase 1 (SpI) ^22^. In the case of *A. baumannii*, HphA contained a canonical lipobox motif yet was predicted to be processed by signal peptidase I which disregards the typical lipoprotein processing pathway (Figure 1D).

**Figure 1.**
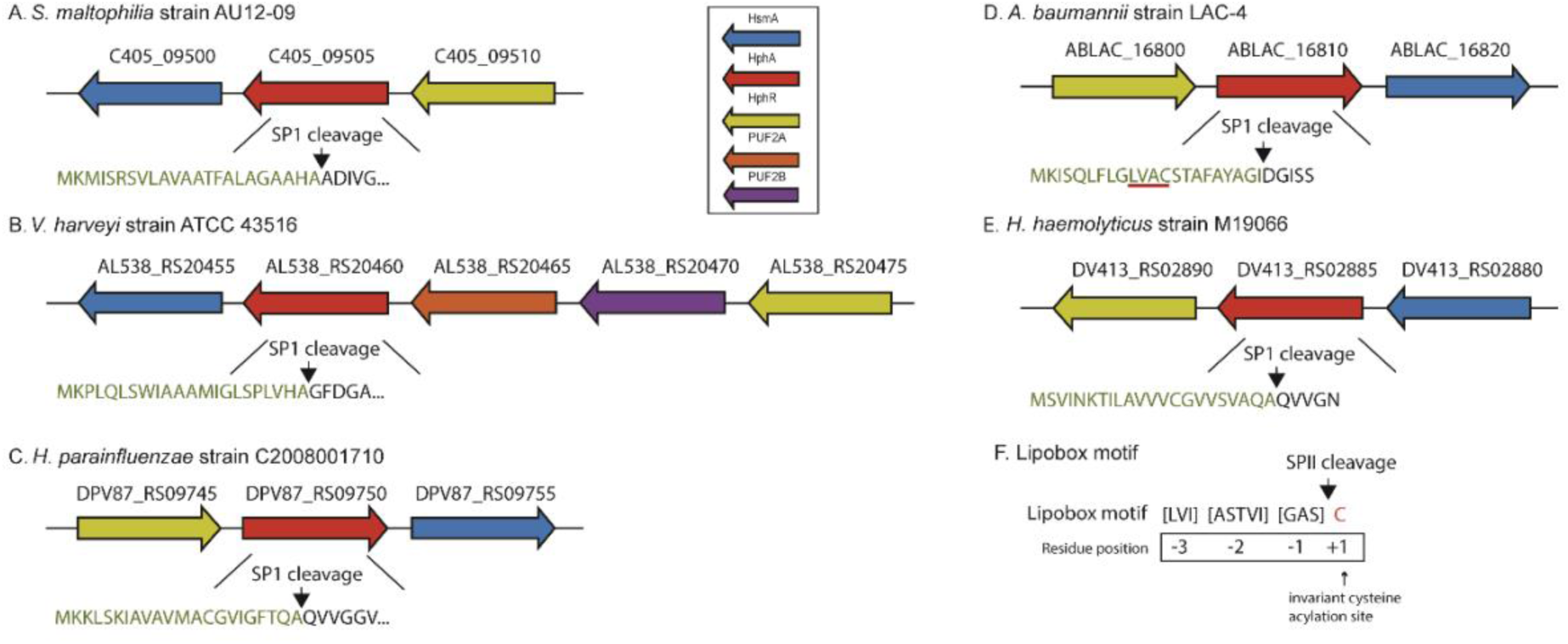
Genomic layout of HphA homolog gene cluster of (A) *S. maltophilia*, (B) *V. harveyi*, (C) *H. parainfluenzae*, (D) *A. baumannii*, (E) *H. haemolyticus*. All HphA gene clusters contain a gene encoding Slam (HsmA), hemophilin (HphA), and TonB-dependent receptor (HphR) which are depicted in blue, red, and green, respectively. An exception is seen in *V. harveyi* where there are two additional proteins of unknown function (PUF) 2A and 2B represented in orange and purple, respectively. The annotation above the arrows represents the NCBI ‘locus tag’ of each gene. The green coloured protein sequence highlights the signal peptide cleaved by signal peptidase I (SpI). The first amino acid residue after the signal peptide cleavage becomes the residue position +1. The putative lipobox motif of bacterial lipoproteins consist of four amino acid residues typically consisting of those amino acids shown in brackets (F). The residue position at +1 is an invariant cysteine residue (coloured in red) that is crucial for acylation.

The structure of *A. baumannii* HphA contained an eight stranded β-barrel domain that is commonly found in Slam-dependent lipoproteins and as expected, was shown to be secreted by Slam^19^. A study from Latham et al. (2019), that reported the first HphA-type hemophore (called hemophilin) in *Haemophilus haemolyticus,* also showed structural similarity to existing lipoproteins. Interestingly, its signal sequence was predicted to be processed by signal peptidase 1, consistent with the absence of a putative lipobox motif^1^. We found that *H. haemolyticus* HphA was also adjacent to genes encoding a Slam and a TBDR (Figure 1E). This raised the question of whether HphA homologs secreted by Slam are prevalent in Gram-negative bacteria as Slam homologs can be found in diverse proteobacteria.

We investigated HphA homologs in *Stenotrophomonas maltophilia*, *Haemophilus parainfluenzae*, and *Vibrio harveyi* which are part of the gammaproteobacteria representing the largest and the most diverse class, containing approximately 250 genera^23^. The HphA homologs in *S. maltophilia*, *V. harveyi*, and *H. parainfluenzae* shared low sequence identity, ranging from 28-39%, to the HphA found in A. *baumannii* (Supplementary Table 1). We found that HphA in these organisms were also located in the Slam gene cluster together with a TBDR, molecular weight ranging from ∼112 to 120 kDa (Figure 1A-C). The genomic layout in *V. harveyi* differed from the other organisms in that it contained two additional genes of unknown function (PUF2A and PUF2B) between the Slam and TBDR gene (Figure 1B). PUF2A and PUF2B were predicted to have signal peptides for Sec-dependent secretion and contained lipobox motifs however, their relation to Slam is yet to be tested. The signal peptide of all HphA homologs were predicted to be processed by signal peptidase I, suggesting that these are non-lipidated Slam substrates.

### Structural diversity of HphAs stems from the N-terminal heme binding domain

We structurally characterized the HphA homologs with relatively low sequence identity to *A. baumannii* and confirmed their function as heme-binding proteins. The proteins were recombinantly expressed and co-purified with heme in *E. coli*. The crystallized HphA proteins from *S. maltophilia, V. harveyi,* and *H. parainfluenzae* were then subjected to X-ray for structure determination and were solved at the resolution of 1.80 Å, 1.73 Å, and 1.9 Å, respectively (Figure 2, Supplementary Table 2-4). Despite low sequence identity, the newly solved structures of HphA were structurally similar to HphA structures in *A. baumannii* and *H. haemolyticus*. The HphAs all share a C-terminal eight stranded β-barrel paired with an N-terminal β-handle domain. These structural features are also seen in Neisserial surface lipoproteins such as Tbpb, LpbB, HpuA, fHbp, and NHBA (Supplementary Figure 1). Although these lipoproteins and HphAs share the eight stranded β-barrel architecture, each has acquired a different function through their β-handle domain that is beneficial for bacterial survival in the host. Furthermore, the main structural differences between the HphAs are located in the N-terminal domain that utilize unique structural elements to clamp the bound heme group.

**Figure 2.**
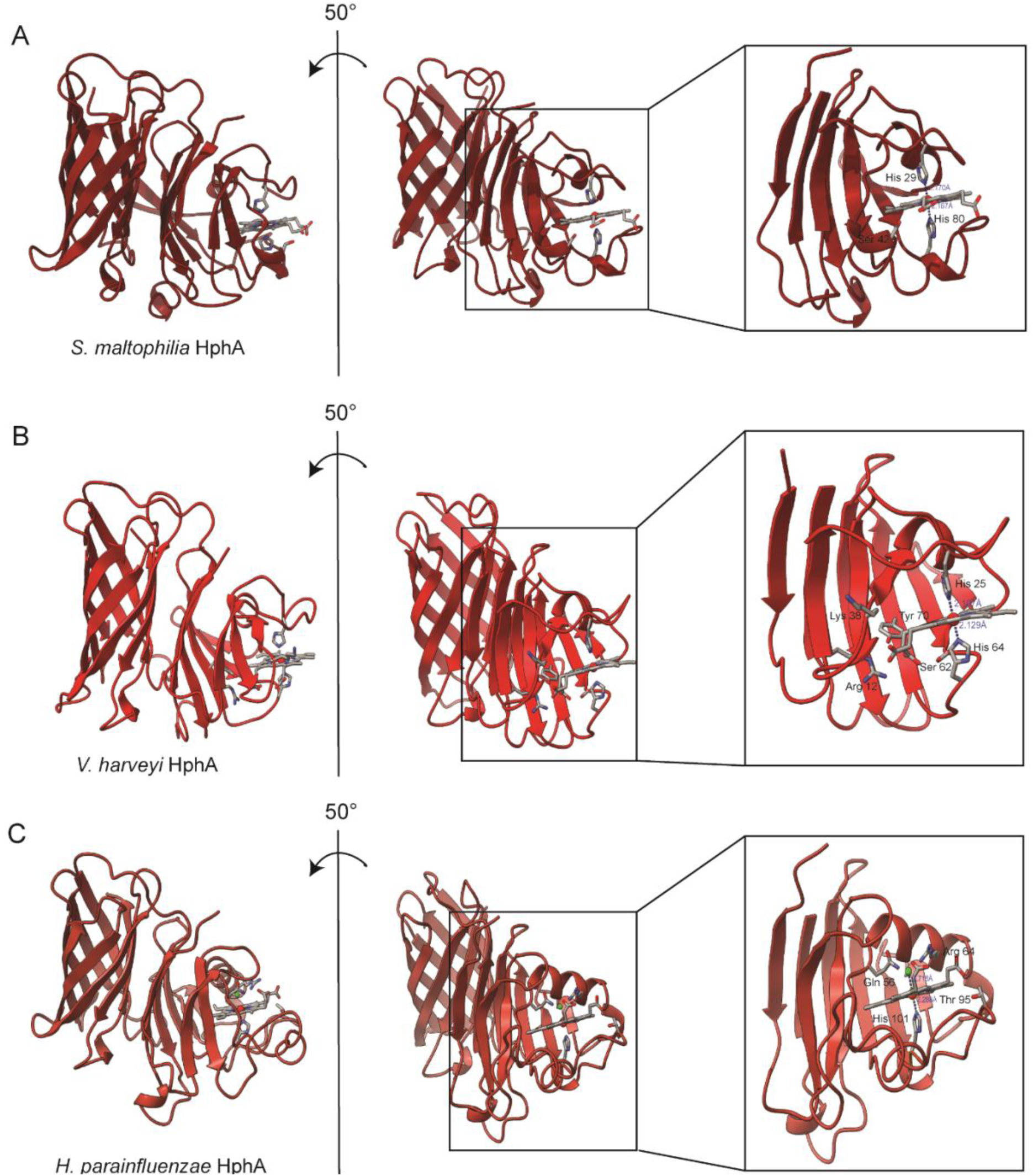
The structure of HphAs in *S. maltophilia*, *V.harveyi*, and *H.parainfluenzae* are shown. Each structure is rotated 50° based on the y-axis. The boxed figure on the far right of each structure is a detailed view of the heme-binding site. Residue side chains that are interacting with the heme are shown in stick figures. The distances of the heme axial ligand are shown in blue. Figures were generated using ChimeraX (Version 1.3). (A) In *S. maltophilia* His29 and His80 acts as heme axial ligands while Ser42 forms hydrogen bond with the propionate group of heme. (B) In *V. harveyi*, His25 and His64 acts as heme axial ligands while Arg 12, Lys 38, Ser 62, and Tyr70 forms hydrogen bonds with the propionate groups of the heme. (C) In *H. parainfluenzae*, His101 acts as the single heme axial ligand while Gln56, Arg 64, Thr95 form hydrogen bonds with the propionate groups of the heme. The distances between the His (NE2) and Fe are indicated in blue. The distance between the chloride ion (lime green) and Fe (red) is also indicated in blue in the *H. parainfluenzae* HphA structure.

A comparison of the five HphA (hemophilin) structures, three determine herein, revealed differences in heme coordination and surface properties. In *S. maltophilia,* the heme is coordinated by two histidine residues, His29 and His80 (Figure 2A). The porphyrin ring is oriented in the heme binding site where the propionate side chains face outwards towards the solvent. One of the propionate groups forms hydrogen bonds (H-bonds) with Ser42. This is similar to heme coordination in *A. baumannii* where His22 and His85 coordinates heme while Tyr38 and Ser84 form polar contacts with one of the propionate groups (Supplementary Figure 2).

Similarly, *V. harveyi* utilizes two histidines (His25 and His64) as heme axial ligands. However, the propionate side chains of the porphyrin ring are facing the loops between the β-strands of the N-terminal domain rather than being exposed to the solvent. Also, polar contacts with the porphyrin ring were observed the most in *V. harveyi*. One of the propionate groups forms H-bonds with Arg12, Ser62, and Tyr70, while the other propionate group forms an H-bond with Lys38.

On the other hand, heme in *H. parainfluenzae* is coordinated by a single histidine residue, His101, and the propionate groups of the porphyrin ring form hydrogen bonds with Gln56, Arg64 and Thr95 residues. This is also observed in the other *Haemophilus* species, *H. haemolyticus* which shares 66% sequence identity with *H. parainfluenzae* HphA based on the aligned 277 amino acid sequences (Supplementary Table 1). In *H. haemolyticus*, His97 acts as a single heme axial ligand and Gln52, Arg60, Thr91 forms polar contacts with the porphyrin ring.

The histidine residue in the loops between β5 and β6 strands are observed to be conserved in *Haemophilus* sp. and *S. maltophilia* HphAs. On the other hand, *V. harveyi* and *A. baumannii* contains an extra, short β-strand consisted of three amino acid residues after β3 strand. Therefore, one of the histidine residues that coordinates the heme in *V. harveyi* is situated between the loops between β6 and β7 strands instead. The heme axial ligand, histidine, in *A. baumannii* is located on a short helix that is also situated between the β6 and β7 strands. The additional heme axial ligand (histidine residue) seen *in S. maltophilia*, *V. harveyi* and *A. baumannii* is found in the loops between the β2 and β3 strands. The loop between β2 and β3 strands in the *Haemophilus* species is oriented away from the heme while the same loop in *S. maltophilia*, *V. harveyi*, and *A. baumannii* is directed towards the heme. These differences in heme coordination may reflect difference in how tightly each HphA binds to the heme ligand. Since some bacterial species colonize the same niche, it is possible that the bacteria have evolved to compete for the same source of heme by presenting a more effective heme binding domain.

### HphAs are present in alpha-, beta- and gamma-proteobacteria

With the independent discoveries of HphA in *A. baumannii* and H. *haemolyticus* HphA, we next conducted an exhaustive investigation into the diversity and prevalence of the Slam-dependent hemophore system in Gram-negative bacteria by bioinformatic analysis. Five sequences corresponding to the solved hemophore structures including the three new structures reported in this study were queried in a BLASTp search in order to generate an expansive list of putative HphAs. Putative HphAs were narrowed down by filtering for sequences that contained a prevalent Motif-1 (QVGTQDVYFGEWS) obtained from MEME analysis using 540 unique accessions using strict criteria as mentioned in the methods. This filtering process generated the ‘lenient set’ which contained 1550 unique accessions in total (Supplementary Data 1). From this analysis, 76, 276, and 1191 HphA homologs were found to be classified as alpha-, beta-, and gamma-proteobacteria, respectively (Figure 3). These homologs belonged to the following orders: Aeromonadales (2), Alteromonadales (9), Burkholderiales (68), Caulobacterales (10), Chromatiales (1), Enterobacterales (413), Hyphomicrobiales (3), Moraxellales (243), Neisseriales (197), Nitrosomonadales (2), Oceanospirillales (31), Pasteurellales (85), Pseudomonadales (110), Rhodocyclales (7), Sphingomonadales (63), Thiotrichales (9), Vibrionales (58), Xanthomonadales (228) and 11 sequences with unresolved lineages. The bioinformatic analysis showed that the HsmA-HphA-HphR gene cluster is prevalent in diverse Gram-negative bacteria. HphA homologs were not identified in the delta- or epsilon- proteobacteria. Previous Slam bioinformatics analysis indicated few Slam homologs in delta- and epsilon- proteobacteria which suggests why Slam-dependent HphA homologs were absent in these classes.

**Figure 3.**
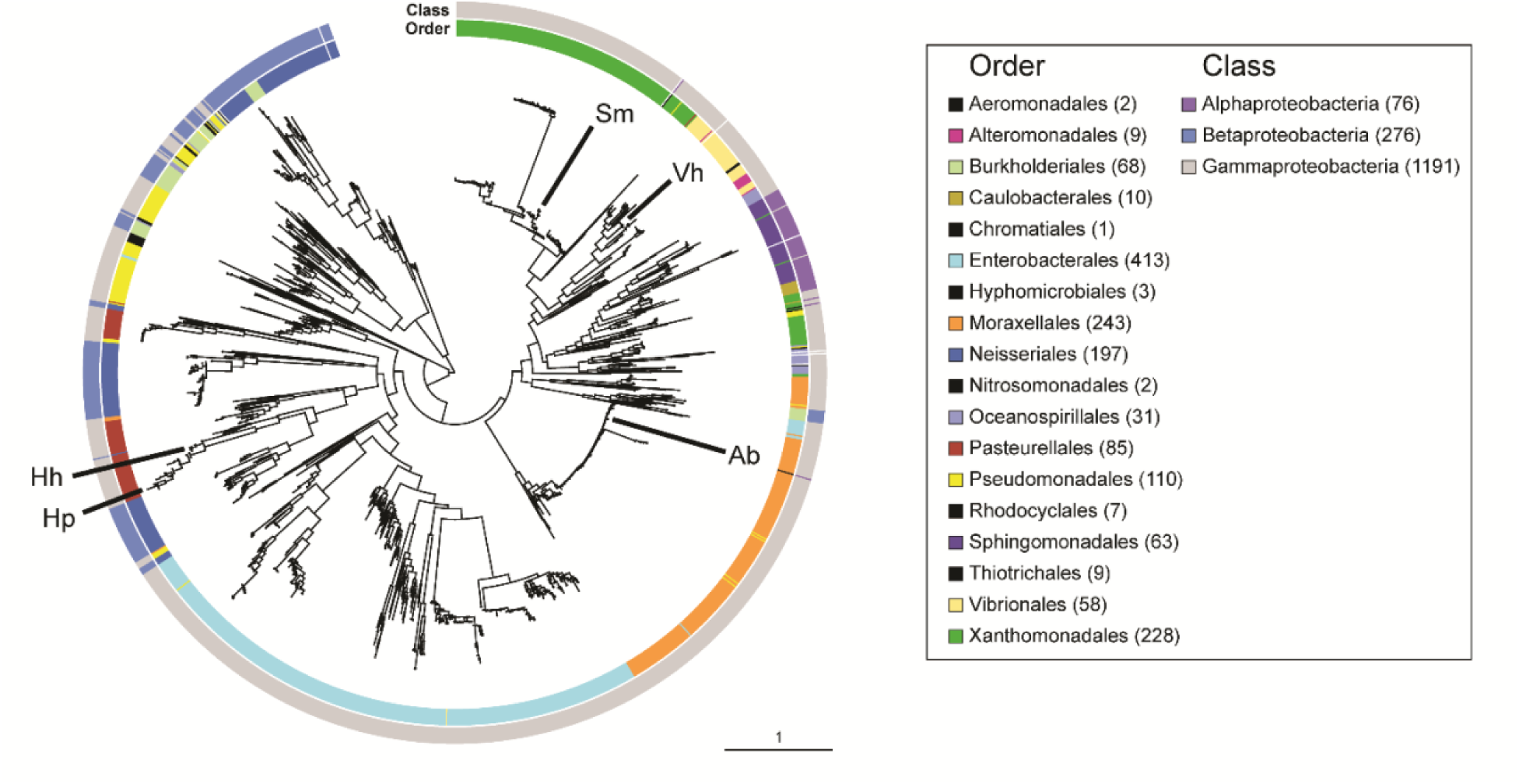
Homologs of HphAs in Proteobacteria from the ‘lenient set’. The lenient set contains a total of 1550 unique sequences. The inner banner indicates the order while the outer banner surrounding the tree represents the class. Five structurally identified HphAs are indicated in the arrows. The annotation Ab, Hp, Hh, Vh, Sm indicates *A. baumannii*, *H. parainfluenzae*, *H. haemolyticus*, *V. harveyi*, and *S. maltophilia*, respectively. The numbers inside the brackets indicates the number of unique accessions found in each class or order. The trees were generated using RAxML.

To identify HphA homologs with higher confidence, we further filtered the ‘lenient set’ for the presence of Motif-3 (MPPSHSALGNFNF) and Motif-9 (AFVDFSGLKGYAQ). These motifs either contained the histidine coordinating the heme ligand (Motif-3) or residues that made polar contacts with the porphyrin ring (Motif-9), yielding 1212 sequences. We then performed a high- throughput genomic context screen to select for sequences that contained a Slam and TBDR genes adjacent or nearby the putative HphA homolog. This analysis yielded in 1015 sequences of HphA homologs with higher confidence comprising the ‘strict tree’ (Supplementary Figure 3, Supplementary Data 2). Homologs in the ‘strict tree’ belonged to the following orders: Aeromonadales (2), Alteromonadales (8), Burkholderiales (10), Caulobacterales (8), Chromatiales (1), Enterobacterales (309), Moraxellales (206), Neisseriales (38), Nitrosomonadales (2), Oceanospirillales (21), Pasteurellales (54), Pseudomonadales (45) Sphingomonadales (39), Thiotrichales (9), Vibrionales (54), Xanthomonadales (205) and 3 sequences with unresolved lineages. This filtering process resulted in a massive reduction in the gamma- and beta- proteobacteria class and mostly from the Enterobacterales and Neisseriales.

### Secretion of HphA is dependent on the presence of Slam

The genetic organization of HphA contains Slam and TBDR as neighbouring genes. Given the past literature on Slam about its crucial role in translocating lipoproteins to the cell surface and that Slam-dependent lipoproteins contain an eight-stranded β-barrel domain, we wanted to test if HphA secretion is dependent on Slam^12, 20^. To distinguish secreted protein-related Slams from lipidated-protein Slams, we renamed this HphA-related Slam as HsmA (<u>h</u>emophilin <u>s</u>ecretion <u>m</u>odulator <u>A</u>)^19^. Consequently, the TBDR is renamed as HphR^19^. The secretion assay was conducted in *E. coli* C43(DE3) or *E. coli K-12* Δ*degP* cells containing plasmids expressing HsmA and HphA from *H. haemolyticus*, *V. harveyi*, and *S. maltophilia*. The cells were grown in autoinduction media to express HsmA overnight while HphA, under the pBAD promoter, stayed uninduced. The next day, cells were diluted into minimal media M63 to OD600 = 0.2 and were induced with 0.5% L-arabinose for HphA expression. The supernatant and whole cell lysate samples were collected at different time points. Western blot analysis was performed to track secretion of Flag-tagged HphA and to ensure the expression of His-tagged HsmA (Figure 4A). Increased secretion of Flag-tagged *H. haemolyticus* HphA was seen in HsmA expressing cells. Minimal secretion of HphA was observed in cells containing an empty vector and Flag-tagged HphA. Therefore, the results showed that the efficiency of HphA secretion is significantly improved by the presence of HsmA. Interestingly, at 22 hr post-induction of *H. haemolyticus* HphA, we observed secretion of HsmA in the supernatant. This was suggested to be HsmA secreted in outer membrane vesicles. In the study of Grossman et al. (2021), a similar observation was shown in *Xenorhabdus nematophilia* where HrpB (HsmA homolog) was detected in the supernatant. However, the ultracentrifugation of the supernatant removed HrpB in the media, suggesting its localization in outer membrane vesicles.

**Figure 4.**
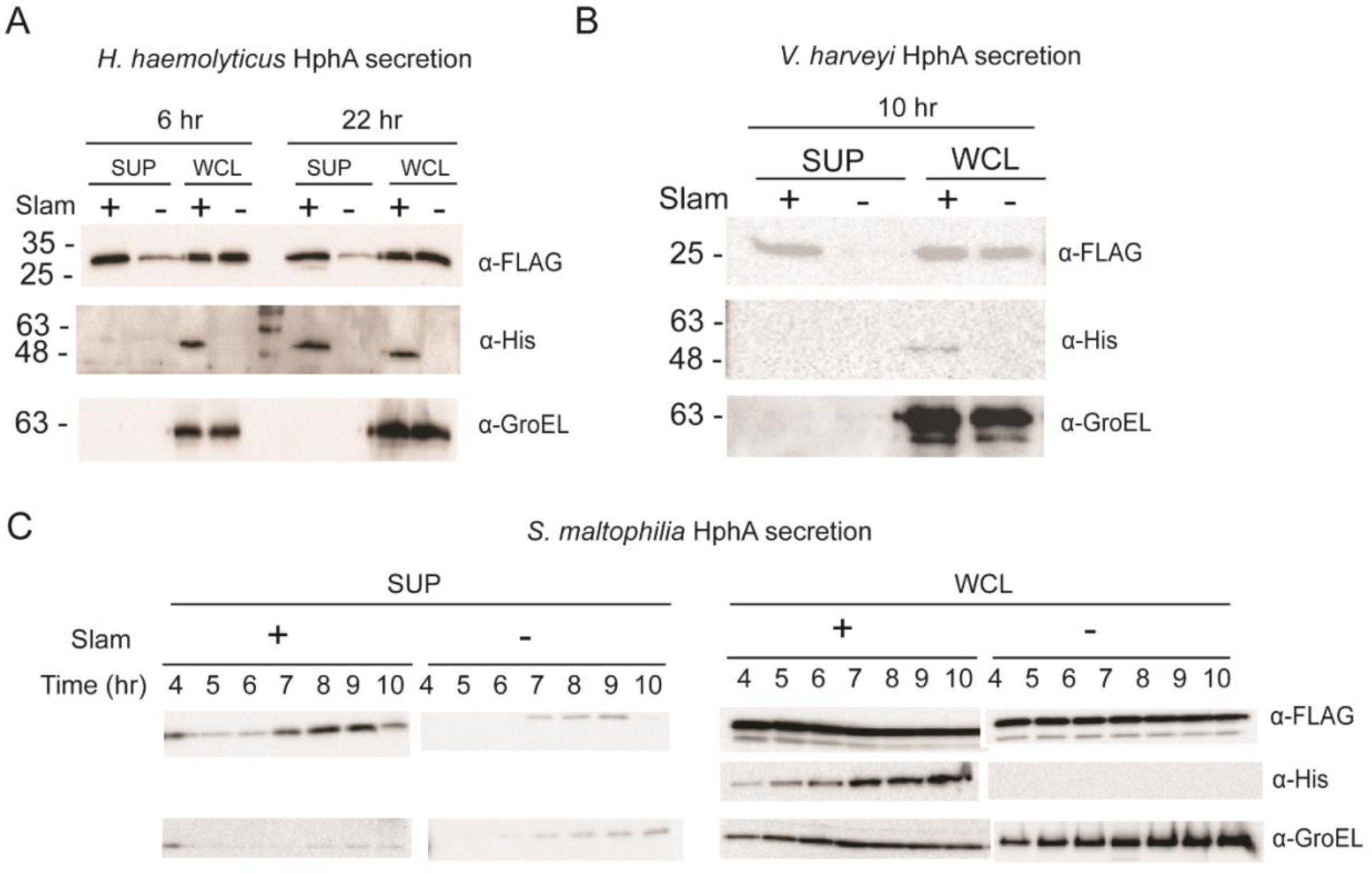
Reconstitution of (A) *H. haemolyticus*, (B) *V. harveyi*, and (C) *S. maltophilia* HphA secretion in *E. coli* cells. (A) FLAG-tagged *H. haemolyticus* HphA with the endogenous signal peptide was expressed alone or with His-tagged HsmA in E. coli C43 cells grown in M63 media. The supernatant (SUP) and the whole cell lysate (WCL) at 6 hr and 12 hr post HphA induction were analyzed by immunoblot probed with α-FLAG, α-GroEL, and α-His to detect the localization of HphA, GroEL, and HsmA, respectively. GroEL is a cytoplasmic chaperone and was used as a cell lysis control. (B) FLAG-tagged *V. harveyi* HphA with the endogenous signal peptide was expressed alone or with His-tagged HsmA in *E. coli K-12 ΔdegP* cells grown in M63 media. The supernatant and the whole cell lysate were analyzed post 10 hr induction of HphA by immunoblot probed with α-FLAG, α-GroEL, and α-His. (C) FLAG-tagged *S. maltophilia* HphA with the endogenous signal peptide was expressed alone or with His-tagged HsmA in E. coli C43 cells grown in M63 media. The supernatants and the whole cell lysate were collected hourly from 4 hr to 10 hr post induction of HphA. Again, the expression of HphA, HsmA, GroEL were assessed by western blot analysis.

In the *V. harveyi* HphA secretion assay, we observed that HphA expression in the whole cell lysate samples without HsmA was significantly lower than whole cell lysates with HsmA when expressed in *E. coli* C43 (DE3) cells. To assess whether DegP was involved in the removal of accumulated *V. harveyi* HphA in the periplasm, we repeated the secretion assay in *E. coli* K12 Δ*degP* strain instead (Figure 4B). We observed that HphA showed similar expression levels in cells with or without HsmA, inferring that DegP is involved in the quality control of HphA that accumulates in the periplasm. Moreover, we detected HphA only in the supernatant from cells expressing HsmA, suggesting that its secretion is dependent on Slam. Lastly, the secretion of *S. maltophilia* HphA was assessed in *E. coli* C43 (DE3) cells. Secretion of HphA was detected post 4 hour of induction and was shown to have increasing levels of HphAs between post 7 to 9 hr. Some cell lysis was observed in both Slam absent and present samples. However, increased HphA secretion was only observed in cells expressing Slam. In conclusion, all HphAs next to the Slam homolog, HsmA, were shown to be secreted into the media when Slam is expressed.

### TonB-dependent receptor confers specificity to its cognate HphA

Since HphA homologs are found in several bacterial species, we wanted to see if HphAs from different species can be shared to support the growth of *A. baumannii*. We performed growth assays in *A. baumannii* with or without its TonB-dependent receptor, HphR, and used HphAs from *A. baumannii*, *V.harveyi*, and *S. maltophilia* as sole heme sources (Figure 5A). In the growth assay, we observed that *A.baumannii* can utilize its cognate HphA most efficiently. However, *A. baumannii* was also able to use *V. harveyi* HphA to some extent while *S. maltophilia* HphA did not support growth. When TonB dependent receptor, HphR, is knocked out in *A. baumannii*, the bacteria can no longer import heme and therefore, growth is suppressed severely (Figure 5B). Together, these results led to a conclusion that TBDR is important for heme uptake and that it confers specificity to its cognate HphA.

**Figure 5.**
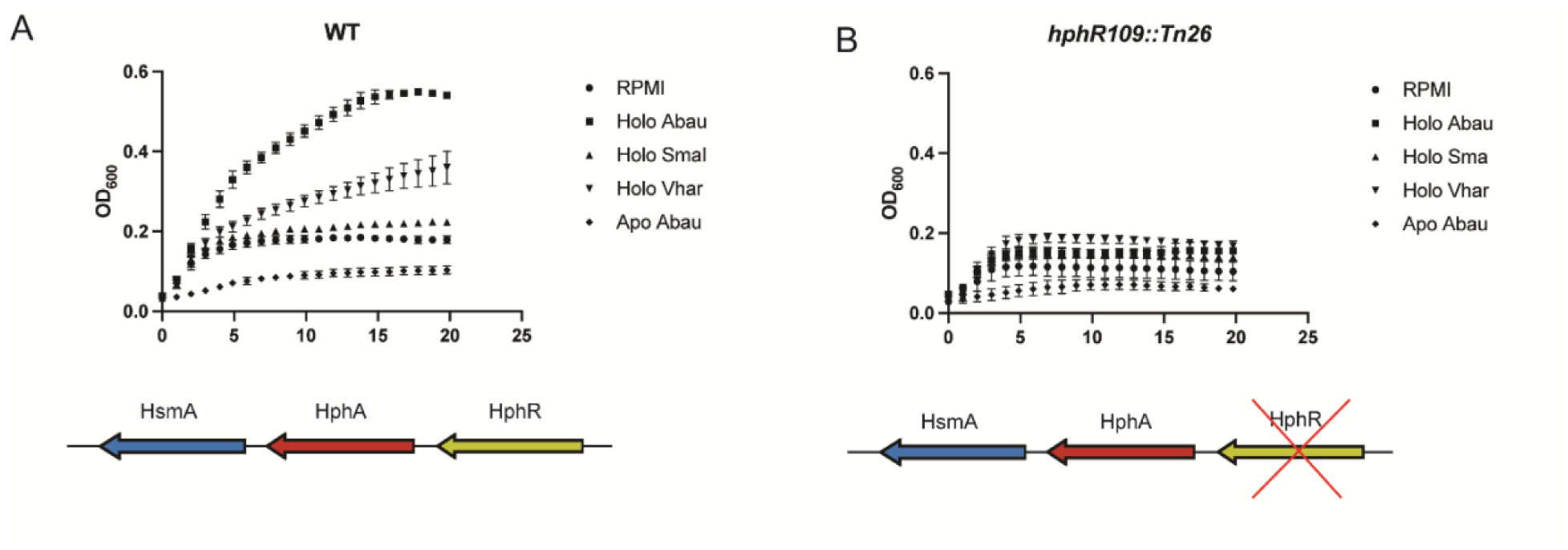
*A. baumannii* growth assays using HphAs from different bacterial species. HphA homologs differ in their ability to support *A. baumannii* growth. To examine specificity in the HphA-R interaction, HphA homologs from *A. baumannii* (Abau), *S. maltophilia* (Smal) and *Vibrio harveyi* (Vhar) were tested for their ability to promote (A)WT and (B) HphR mutant *A. baumannii* growth as the sole iron/heme source in iron limiting media. Plotted data points represent the mean ± SEM for three independent experiments.

## Discussion

In this study, we focused on a heme acquisition system by a novel hemophore called HphA. Here we reported additional structures of HphA hemophores in *S. maltophilia*, *V. harveyi*, and *H. parainfluenzae*. Bioinformatic analysis revealed that the Slam-HphA-TBDR gene cluster is found in many Gram-negative bacteria, restricted to alpha-, beta-, and gamma-proteobacteria. We confirmed that all three genes in this cluster work together to acquire heme. HphA secretion is mediated by the Slam homolog, HsmA, and the import of heme across the outer membrane was fulfilled by the tonB-dependent receptor (TBDR), HphR. The repertoire of Slam substrates has expanded to now include both lipidated and non-lipidated proteins. This study is in agreement with the bioinformatic analysis of Slam homologs performed by Grossman et al. (2021) where the authors found Slam homologs adjacent to genes encoding a non-lipidated substrate that contains the Pfam domain, TbpB_B_D, present in known Slam-dependent substrates^22^.

While numerous heme acquisition methods through surface proteins have been reported, secretion of hemophores is another powerful strategy employed by many Gram-negative bacteria to maximize efficiency to acquire heme by overcoming diffusion limitations^24^. The working model of heme acquisition by HphA is as follows (Figure 6): the HphA nascent chain is synthesized in the cytoplasm with its signal peptide. HphA is then translocated across the inner membrane by the Sec pathway. Although HphA structurally resembles Slam-dependent surface lipoproteins that contains the eight-stranded β-barrel domain, the protein sequence does not contain a putative lipobox motif and therefore, is predicted to be processed by the signal peptidase I. HphA is then suggested to interact with periplasmic chaperone(s) that keep the protein unfolded in the periplasmic space. The HsmA (Slam homolog) then translocates HphA across the outer membrane. The properly folded apo-HphA then carries on its role to scavenge free heme from the host environment. Holo- or heme loaded HphA recovers heme via the outer membrane TonB-dependent receptor (TBDR), HphR (As shown by the alpha-fold predicted complex of *S. maltophilia* HphA and HphR). The inner membrane TonB complex interacts with the TBDR to import heme into the periplasm. Subsequently, periplasmic heme binding protein shuttles heme to the ABC transporter in the inner membrane to complete its import into the cytoplasm. Heme can then bind to cytoplasmic hemin-binding protein and be degraded by heme oxygenase.

**Figure 6.**
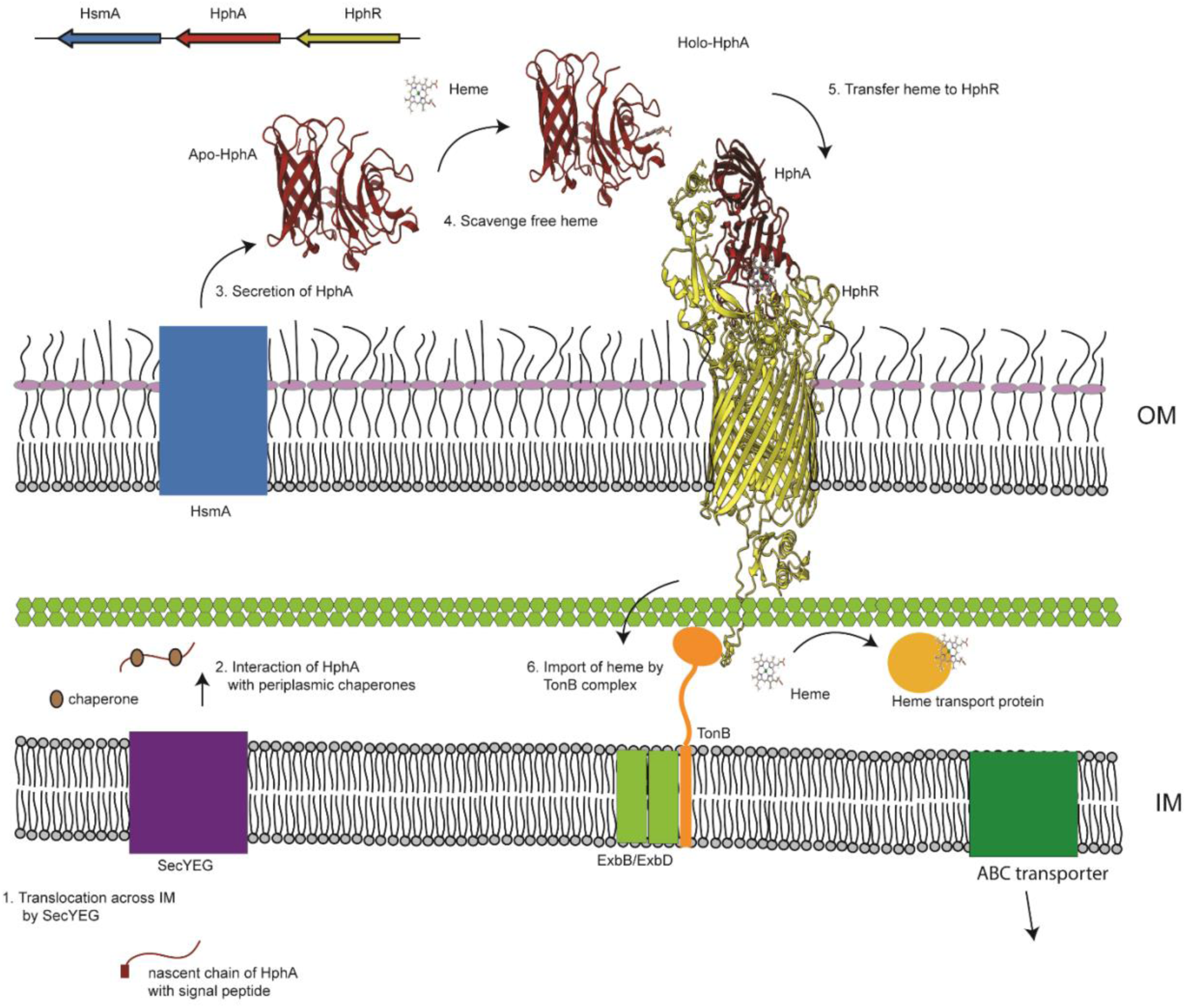
Working model of heme acquisition by HphA. The nascent chain of HphA with its signal peptide gets translocated across the inner membrane (IM) by the SecYEG translocon followed by the signal peptidase I (SpI) cleaving its signal peptide. The HphA is shuttled across the periplasm through the help of chaperones that delivers it to HsmA (Slam homolog). The HsmA then translocates HphA across the outer membrane (OM) and into the extracellular environment where the HphA scavenges the heme from the host environment. Heme-loaded HphA then interacts with the Ton-B dependent receptor, HphR which subsequently triggers the interaction with the TonB complex (TonB-ExbB-ExbD) that uses the energy (coupled to the proton motive force) to import heme across the OM. The AlphaFold2 prediction of HphA:HphR complex is shown in the figure^56^. The heme transport protein in the periplasm shuttles the heme to the ABC transporter which ultimately imports the heme to the cytoplasm.

The structure of HphAs are quite different from existing Gram-negative hemophores such as HasA, HusA, HmuY, and HxuA (Supplementary Figure 5). HasA is a 19 kDa hemophore that was first identified in *S. marcescens* and was shown to use Type 1 secretion system for its secretion^25^. The structure of HasA consist of four α-helices and seven β-strands and contains two axial heme ligands His32 and Tyr75 coordinating heme that binds at Kd 18 pM^14, 26–29^. HusA consist of nine α-helices forming a right-handed super-helical arrangement and binds heme on the concave surface at Kd of µM range^17^. HmuY is an all β-fold protein, that resembles a right hand, consisting of 15 β-strands binding the heme between the beta strands mimicking the thumb and ring-finger with low affinity (Kd∼ 3µM)^16^. Lastly, HxuA is a 100 kDa unconventional hemophore that was shown to be bind hemopexin rather than heme^30, 31^. HxuA is consist of two domains; N-terminal secretion domain, also known as the TPS domain, and the C-terminal functional domain which together is organized into a right-handed β-helix^32^. While most HxuA was appeared to be anchored, a small proportion of functional HxuA was found to be secreted to the extracellular medium in the reconstituted *E. coli* system^31^. On the other hand, HphA consists of an N-terminal clamp-like domain that binds to heme and a C-terminal eight-stranded β-barrel domain which resembles β-barrel domains of Slam-dependent lipoproteins (Supplementary Figure 1). The function of this β-barrel domain is unknown but it has been suggested to be important for secretion by Slam^33^. In conclusion, HphA represents a new structural class of hemophores as its overall structural fold and the heme binding domains differ from the existing Gram-negative hemophores.

Given that the most abundant heme source is hemoglobin, it is likely that HphA obtains its heme from heme sources such as hemoglobin. As free heme in not readily available to invading pathogens, certain bacteria can secrete hemolysin or hemoglobin proteases in order to release free hemin from heme loaded host proteins^34, 35^. Apart from hemoglobin, hemoglobin-haptoglobin complex, hemopexin, myoglobin, albumin, and cytochromes can also serve as heme sources for bacteria^36^. The ability to acquire heme is important for heme auxotrophs as heme not only acts as an iron source but also a protoporphyrin source for organisms that are unable to synthesize the tetrapyrrole ring. For instance, *H. haemolyticus* and *H. influenzae* are unable to synthesize their own heme so having such heme acquisition systems as HphA would benefit these organisms. Interestingly, *H. parainfluenzae* is capable of synthesizing its own heme and a smaller percentage of *H. parainfluenzae* also contains HphA, which is the protein structure we solved^1, 37^. In order to compete and colonize in the same niche as the other Haemophilus species, they may have acquired this system over the course of evolution.

This HphA system is also found in *S. maltophilia* which is capable of synthesizing heme. The iron/heme acquisition system in *S. maltophilia* have not been well characterized. In silico analysis have revealed putative iron and heme acquisition systems in *S. maltophilia*^38^. A recent study elucidated the first hemin acquisition system in *S. maltophilia* by a TonB-dependent receptor called HemA which was shown to be a similar homolog of PhuR from *Pseudomonas aeruginosa*^39^. In this paper, we report the first hemophore in *S. maltophilia*. The Slam-dependent HphA hemophore gene cluster can be found in 139 out of 141 *S. maltophilia* genomes suggesting their importance of HphA in heme acquisition (based on tBLASTn search, with a threshold of 1E-50, using 10 representative sequences Besides the colonization in humans, S. maltophilia has been reported to colonize and infect in a wide range of animals, including mammals, fish, reptiles and particularly in horses^40^. This suggests that HphA can utilize heme sources from diverse organisms. Apart from its role in effectively scavenging heme for growth in different organisms, HphA was shown to partake in another interesting role. The HphA from *S. maltophilia* K279a, denoted as *smlt2713* or SMLT_RS12935 was also shown to be secreted into the media^41^. SMLT_RS12935 shares 86.61% protein sequence identity with the HphA in AU12- 09 strain. The authors discovered this protein while exploring microbial signals that regulate the anti-inflammatory characteristics of gut-resident macrophages^41^. They found that *S. maltophilia* is a persistent colonizer within non-lymphoid mucosal tissue-resident macrophages and helps to maintain homeostatic condition in the colon through the induction of IL-10 which is enhanced by *smlt2713*^41^. The absence of smlt2713 or deficiency of IL-10 impaired the ability of *S. maltophilia* to intracellularly colonize macrophages^42^. Moreover, in a transcriptomic analysis of *S. maltophilia* under biofilm conditions showed that HphA was the most strongly and differentially expressed gene in all strains^43^.

The mechanism of heme stealing by HphA is not known. However, in our previous studies, it was suggested that the heme binding of HphA is by a passive process rather than an active pirating process as the Soret signature of a known heme scavenger hemopexin showed similar levels to HphA after incubation with the hemoglobin resin^19^. Heme could be passively transferred from a lower affinity HphA to a higher affinity TBDR or in the case of the HasA system, the lower affinity HasA receptor (HasR) actively opens HasA using one of its extracellular loops^28^. The overall structure between the structurally solved HphAs were similar (Supplementary Figure 6). While all HphAs bound to heme through the N-terminal domain, subtle differences in heme coordination (single vs two histidine residues) and porphyrin interacting residues were observed. These differences may result in different affinities to bind heme and its ability to scavenge heme from host hemoproteins. Many of these bacterial species often colonize in the same niche. For instance, *S. maltophilia* and Haemophilus sp. are both commonly found in respiratory tracts^44^. It is possible that some species may have evolved to contain a tighter binding mechanism to compete with HphAs produced by a different organism. Moreover, our analysis of hemophilin gene clusters in multiple bacteria demonstrate the specificity in this system (Figure 5A, B). Considering the overlap in host niches for some of these bacteria highlighted in this study, the secretion of HphA for host heme scavenging is beneficial in a competitive landscape where commensals and even other opportunistic pathogens are vying for the same resources. Future studies characterizing the heme binding affinity for the different HphA homologs will also have important implications on the mechanism of heme transfer. In addition, specificity of the hemophilin systems can be exploited for the targeted delivery of therapeutics to pathogens leaving the natural microbiome unperturbed. Heme analogs that contain a redox inactive gallium instead of iron are already being developed and tested, showing promise for combating multi-drug resistant pathogens^45^.

## Materials and methods

### Bacterial strains, plasmids, primers, and antibodies

Please refer to supplementary Table 1 for a summary of bacterial strains, plasmids, primers and antibodies used in this study.

### Plasmid construction and cloning of HphAs and HsmAs

For constructs used for purification and crystallization studies, synthetic DNA sequences of gene locus tag AL538_RS20460 from *V. harveyi* and C405_09505 from *S. maltophilia*, excluding the endogenous signal peptide region, were cloned into pET52b plasmid with a C-terminal thrombin-cleavable 10x or 6x histidine (His) tag, respectively. Synthetic DNA sequences of gene locus tag WP_111316019 from *H. parainfluenzae* excluding the endogenous signal peptide region were cloned into pET52b vector containing TEV cleavable N-terminal protein sequence that contains 8x His-tag, biotin acceptor peptide (BAP)-tag, and maltose binding protein (MBP) in sequence. Restriction-free (RF) cloning method was used as described in van den Ent & Löwe (2006)^46^.

For constructs used for secretion assays, the HphA locus tags, including the endogenous signal peptide, AL538_RS20460 from *V. harveyi*, C405_09505 from *S. maltophilia*, 6OM5_A from *H. haemolyticus* were cloned into pHERD20T vector with a C-terminal FLAG sequence. The Slam homologs C405_09500 from *S. maltophilia*, WP_050936331.1 from *V. harveyi*, and WP_118817043 from *H. haemolyticus* were cloned into pET26b vector containing a N-terminal 6x His tag and a pelB signal peptide, replacing the endogenous signal peptide. Again, RF cloning method was used as described^46^.

### *S. maltophilia* HphA protein expression and purification

*E. coli* C43(DE3) cells were used to cytoplasmically express *S. maltophilia* HphA (pET52b) in autoinduction media (1% tryptone, 0.5% yeast extract, 0.5% glycerol0.05%, D-glucose, 0.2% α- lactose, 50 mM Na_2_HPO_4_, 50 mM K*H*_2_*PO*_4_, 25 mM (N*H*_4_)_2_S*O*_4_), 2 mM MgS*O*_4_) supplemented with 100 µg/ml ampicillin at 37C for 18 hr^47^. The cells were pelleted and resuspended in lysis buffer [25 mM Hepes pH 7.5, 300 mM NaCl, 10 mM imidazole, lysozyme (1 mg/ml), 2 mM PMSF, 1 mM Benzamidine, DNase I (0.05 mg/ml)] and subsequently lysed using a high pressure homogenizer (Avestin Emulsiflex C3, ATA scientific) at 1000 – 1500 bar pressure. Cell lysate was centrifuged at 17,000 rpm (JA-25, Beckman Coulter) for 1 hr and the supernatant was filtered using 0.45 µm filter before subjecting it to Ni-NTA resin (Thermo) at 4°C overnight. The resin was washed thoroughly with the wash buffer (25 mM Hepes pH 7.5, 300 mM NaCl, 20 mM imidazole) and eluted using 25 mM Hepes pH 7.5, 300 mM NaCl, 250 mM imidazole. The protein was concentrated to a lower volume using 10kDa molecular weight cut-off concentrator (Vivaspin) and injected onto the Superdex™ 75 10/300 GL column equilibrated with 25 mM Hepes pH 7.5 and 100 mM NaCl.

### *V. harveyi* HphA protein expression and purification

*E. coli* Shuffle T7 cells were used to grow transformants containing pET52b *V. harveyi* HphA in LB supplemented with 100 μg/mL ampicillin at 37°C until the *OD*_600_ reached approximately 0.6-0.8. Protein expression was then induced at 1mM IPTG for 18 hr at 20°C. The cells were pelleted and resuspended in lysis buffer containing 50 mM Tris pH 8, 300 mM NaCl, 40 mM imidazole, lysozyme (1 mg/ml), 2 mM PMSF and Dnase I (0.05 mg/ml). The cells were lysed using a high pressure homogenizer as mentioned previously. The supernatant was incubated with Ni-NTA resin (Thermo) at 4°C for 2 hours. The resin was washed and eluted with buffer containing 50 mM Tris pH 8, 300 mM NaCl and 400 mM imidazole. The protein was concentrated to a lower volume using 10kDa molecular weight cut-off concentrator (Vivaspin) and injected onto the Superdex™ 75 10/300 GL column equilibrated with 20 mM Tris pH 8 and 100 mM NaCl.

### *H. parainfluenzae* HphA protein expression and purification

*E. coli* Shuffle T7 cells were used to express *S. maltophilia* HphA (pET52b) in autoinduction media (1% tryptone, 0.5% yeast extract, 0.5% glycerol, 0.05% D-glucose, 0.2% α-lactose, 50 mM Na_2_HPO_4_, 50 mM K*H*_2_*PO*_4_, 25 mM (N*H*_4_)_2_S*O*_4_, 2 mM MgS*O*_4_) supplemented with 100 µg/ml ampicillin at 37 °C for 24 hr^47^. The cells were pelleted and resuspended in lysis buffer [20 mM Hepes pH 7.5, 200 mM NaCl, 10 mM imidazole, lysozyme (1 mg/ml), 2 mM PMSF, 1 mM Benzamidine, Dnase I (0.05 mg/ml)]. Cells were lysed using sonication at 30% duty cycle over the duration of 30 minutes with frequent breaks in between to prevent overheating. The lysate was then centrifuged at 17,000 rpm (JA-25, Beckman Coulter) for 1 hr to remove cell debris. The supernatant was filtered (0.45 µm filter) and repetitively loaded on the HisTrap™ excel 5ml column (Cytiva) overnight at 4 °C using the peristaltic pump (0.5 ml/min). The protein was then eluted using 20 mM Hepes pH 7.5, 200 mM NaCl, 300 mM imidazole. The protein was dialyzed against 20 mM Hepes pH 7.5 buffer at room temperature for 4 hr to reduce imidazole concentration before TEV treatment. Approximately 20 mg of in-house made TEV protease was used via dialysis in 20 mM Hepes pH 7.5, 20 mM NaCl buffer to cleave 320mg of the N-terminal 8x His- and BAP-tag fused MBP protein away from the protein of interest, *H. parainfluenzae* HphA. After TEV cleavage, the negatively charged N-terminal His- and BAP-tag fused MBP protein was separated from HphA by ion-exchange chromatography with HiTrap Q HP 5ml column (GE Healthcare). The flow-through, which contained HphA, was concentrated and further separated from contaminants by size exclusion chromatography using the HiLoad™ 16/60 Superdex™ 75 prep grade column; equilibrated with 20 mM Hepes pH 7.5 and 100 mM NaCl buffer.

### Structure determination of HphAs

Trypsin dissolved in 1x PBS, 2mM Ca*Cl*_2_ was added in 1:100 w/w to purified *S. maltophilia* HphA (32.9 mg/ml) immediately before setting up the sitting drop vapor diffusion in a 1:1 ratio of protein to precipitant for a total of 0.8 µl. HphA was screened against MCSG and JCSG+ commercial screens using a Gryphon robotic drop setter (Art Robbins Instruments). Red crystals were obtained in MCSG1-G7 screen condition: 0.1 M Tris-HCL pH 8.5, 25% (w/v) PEG 3350. Diffraction data were collected at the Canadian Light Source (CLS) in Saskatoon using the ID beamline. The crystals diffracted X-rays to 1.8Å resolution. *A. baumannii* HphA, which shares 39% (alignment length 281) was used as a search model for molecular replacement to solve the phase information of *S. maltophilia* HphA by Phaser in Phenix. The automated and manual refinement was conducted by Phenix and Coot^48, 49^.

*V. harveyi* HphA were also obtained by adding trypsin (1:100 w/w) to the purified HphA (3.35 mg/ml) and was screened against commercial MCSG crystallization screens in 1:1 volume ratio of protein to precipitant for a total of 1 μL. An initial crystal hit was observed in the condition of 25% PEG3350, 0.1 M Bis-Tris pH 5.5. Further optimization of *V. harveyi* HphA crystals was manually performed by altering pH, precipitant concentrations, precipitant compositions, the ratio of protein to precipitant as well as protein concentration. The best crystals were obtained in 25% PEG3350, 0.1 M Bis-Tris pH 5.0 and 20% glycerol with streak seeding and *V. harveyi* HphA at 6.31 mg/ml mixed with trypsin at 0.06 mg/mL. The *V. harveyi* HphA crystals were sent to the Advanced Photon Source (APS) Synchrotron Facility for data collection using the 24ID-C beamline. Datasets of *V. harveyi* HphA crystal diffraction were collected at 0.9791 Å using 0.2° oscillations per image for a total of 1000 images. *S. maltophilia* HphA, which shares 42% sequence identity (alignment length 269) to *V. harveyi* HphA, was used as a search model for molecular replacement to solve the phase information of *V. harveyi* HphA by Phaser in Phenix and was further refined by Phenix and Coot^48, 49^.

Protein crystals of *H. parainfluenzae* HphA was obtained by optimizing crystal growth by microseeding technique from a hit condition that contained red crystals with no defined shape. Using this method, red crystals were obtained in 0.1M Tris-HCl pH 8.52 and 2M ammonium sulfate condition. Diffraction data was collected from National Synchrotron Light Source II (NSLS-II) using FMX beamline. The structure was solved using the first reported hemophilin (or renamed as HphA) in *Haemophilus haemolyticus*, which shares 66% sequence identity with *H. parainfluenzae* HphA based on the aligned 277 amino acid sequences. Crystals were cryoprotected in 20% glycerol.

### LIGPLOT generation

The LIGPLOT program was used to generate a 2-D representation of HphAs and heme ligand interaction in standard PDB data format^50^.

### Identifying putative sequence features of HphA

To search for HphA homologs, we first require ways to identify and define them. We began with the five structurally determined HphAs from *S. maltophilia*, *V. harveyi*, *A. baumannii*, *H. parainfluenzae*, and *H. haemolyticus* as our queries and performed a BLASTp search using an E- value threshold 0.05^51^. The HphAs determined to date were all from the gammaproteobacteria, which is likely to bias the search and miss true divergent hits. To address this, we manually selected from the resulting hits seven additional sequences with low sequence identity (<50%) and repeated the BLASTp search with these twelve queries to ensure we maximized the sensitivity. The seven additional seed sequences were from the following species: *Pseudomonas fluorescens*, *Pasteurella multocida*, *Marinobacterium mangrovicola*, *Xenorhabdus ishibashii*, *Sphingobium baderi*, *Photorhabdus heterorhabditis*, *Neisseria macacae*, and *Neisseria canis*. Filtering out sequences that were >95% identical yielded a total of 1833 hits.

Because we used such a permissive E-value threshold in order to improve the sensitivity of our search at the cost of specificity, we used gene synteny to filter out likely false positive hits. The 1833 hits were analyzed using RODEO2 (http://www.ripp.rodeo/index.html), a tool that can fetch the Pfam profiles of ORFs surrounding a query accession^52^. These hits were filtered for those with neighbors (8 genes upstream or downstream) containing both the Slam and TonB dependent receptor protein domains, resulting in 1078 unique accessions. From this, we further filtered hits for neighbors containing domains relating to iron uptake and processing, including heme oxygenase, heme degradation, heme utilisation, and FecR, which yielded 595 unique accessions. As a final quality control, we performed a multiple sequence alignment and manually removed proteins that did not contain the conserved histidine residue found in the loops between beta strands number 5 and 6, serving as the axial ligand to heme, which ultimately yielded 540 unique accessions that we believe are likely homologs with high confidence.

After removing the predicted signal peptides, these sequences were analyzed for shared motifs using MEME v5.4.1 (Supplementary Figure 4, Supplementary Data 3)^53^. Settings used for MEME analysis were as follow: Motif site distribution: Zero or one site per sequence; Maximum number of motifs: 10; Motif E-value threshold: no limit; Minimum motif width:6; Maximum motif width: 13; Minimum sites per motif: 2; Maximum sites per motif: 540. We identified Motif-1 (QVGTQDVTFGEWS) in all structurally determined HphAs, as well as all highly similar sequences. The iron-coordinating histidine of four of the structurally determined HphAs was located within Motif-3 (MPPSHSALGNFNF) except for the *V. harveyi* HphA. While the histidines residues coordinating the heme iron in *V. harveyi* did not fall into any of the motifs identified in the MEME analysis, Motif-9 (AFVDFSGLKGYAQ) was identified in *V. harveyi* and contained Lys38 which makes polar contacts with the porphyrin ring. This motif was also found in all structurally solved HphA structures except for *H. parainfluenzae*. Together, these data provide ways to identify putative HphAs with varying levels of confidence.

### Identifying and visualizing putative HphA homologs

We supplemented the 1833 possible HphAs with 155 sequences that were located nearby to a Slam subtype we have previously associated with HphAs (Supplementary Data 4), and which contained Pfam domain TbpB_B_D (https://pfam.xfam.org/family/PF01298). This set of 1988 possible HphAs was filtered for those containing Motif-1, due to its ubiquitous presence in our high-confidence set. This yielded 1550 protein sequences that we refer to as the ’lenient set’. These sequences were then filtered for those containing either Motif-3 or Motif-9, which were those identified as structurally important for heme binding, leaving 1212 hits. These were further filtered using RODEO2 to those containing neighbors with both the Slam and TonB dependent receptor protein domains, yielding 1014 sequences. Sequence AIY37036.1, one of the structurally-confirmed HphA homologs, had been removed by RODEO2 likely due to a low-quality genome assembly, so it was manually added to give us a final count of 1015 hits that we refer to as the ’strict set’ (Supplementary Figure 3, Supplementary Data 2).

Both sets of sequences were aligned using MAFFT v7.475, and phylogenetic tree reconstruction was carried out using RAxML v8.2.12 with the PROTGAMMAWAG model of evolution^54, 55^. Trees were visualized and annotated using Navargator (https://compsysbio.org/navargator).

### Slam-dependent HphA secretion assay in *E. coli*

The secretion assay was conducted in *E. coli* C43(DE3) cells containing plasmids expressing pET26 N-terminal 6x His-tag HsmA and pHERD20T HphA with C-terminal FLAG-tag from *V. harveyi, H. haemolyticus* and *S. maltophilia*. The cells were grown in autoinduction media (1% tryptone, 0.5% yeast extract, 0.5% glycerol, 0.05% D-glucose, 0.2% α-lactose, 50 mM Na_2_HPO_4_, 50 mM K*H*_2_*PO*_4_, 25 mM (N*H*_4_)_2_S*O*_4_, 2 mM MgS*O*_4_) supplemented with 100 µg/ml kanamycin and 100 μg/mL ampicillin to express HsmA overnight while HphA, under the pBAD promoter, stayed uninduced. The next day, cells were diluted into minimal media M63 (containing 1mM MgSO4, 0.5% glycerol, 100 μg/mL ampicillin and 50 μg/mL kanamycin) to OD600 = 0.2 and was induced with 0.5% L-arabinose for HphA expression. 2ml of cells were collected at different time points and was centrifuged at 2,348g (FA-45-24-11, Eppendorf rotor) for 5 minutes at 4 °C. The pellet was resupended in 2ml of water and 50 µl of cell resuspension was mixed with 50 µl of 2x SDS loading buffer. The supernatant was centrifuged again at 21,100g (75003424, Thermo Scientific rotor) at 4°C for 15 minute to remove any cell debris and 50 µl of clarified supernatant was added to 50 µl of 2x SDS loading buffer. The cell and supernatant sample collected from different time points were loaded to a 15% SDS-PAGE gel. Semi-dry western blotting was performed using a PVDF membrane. The secretion of HphA, HsmA, and GroEL (loading control) was tracked using α-FLAG (1:5,000), α-His (1: 5,000), and α-GroEL (1:30,000) antibodies, respectively.

### *A. baumannii* growth assays using holo HphAs from *S. maltophilia* and *V. harveyi*

*A. baumannii* strains were iron starved by overnight growth on LB-agar supplemented with 200 µM 2,2’-dipyridyl at 37°C. Cells were resuspended in RPMI, starting at OD_600_ of 0.2, containing 3 µM purified holo HphA from *A. baumannii*, *S. maltophiolia*, and *V. harveyi*. Apo *A. baumannii* HphA was used as a negative control to ensure that the growth is heme-dependent. Apo-HphA was produced by acid-acetone treatment of the holo protein as described in the previous study^19^. Growth at 37°C was monitored over a 20 h period was recorded using a VICTOR Nivo™ plate reader (filter 600/10nm (15mm)).

## Acknowledgements

We would like to thank members of the Moraes, Schryvers and Gray-Owen laboratories for valuable discussions.

## Author Contributions

HES, CP, TJB, and TFM designed experiments and conceived the study. HES, CP, DMC, TJB, DHYC, DN, and MS performed experiments. HES, CP, DMC, TJB, DHYC, DN, MS, and TFM analyzed data. TFM supervised all biochemical work, DMC supervised all bioinformatic analysis. HES, CP, and TJB wrote the manuscript with input from TFM.

## Supporting information

Attached separately

Supplementary Data 1: Fasta file of protein sequences in the ‘lenient set’ of bioinformatic analysis of HphA homologs.

Supplementary Data 2: Fasta file of protein sequences in the ‘stric set’ of bioinformatic analysis of HphA homologs.

Supplementary Data 3: Meme analysis of 540 unique HphA homologs.

Supplementary Data 4: Fasta file of 155 protein sequences that contained the Pfam TbpB_B_D domain next to the Slam subclass that are associated to HphAs from a separate analysis.

---

**Supplementary Figure 1.**
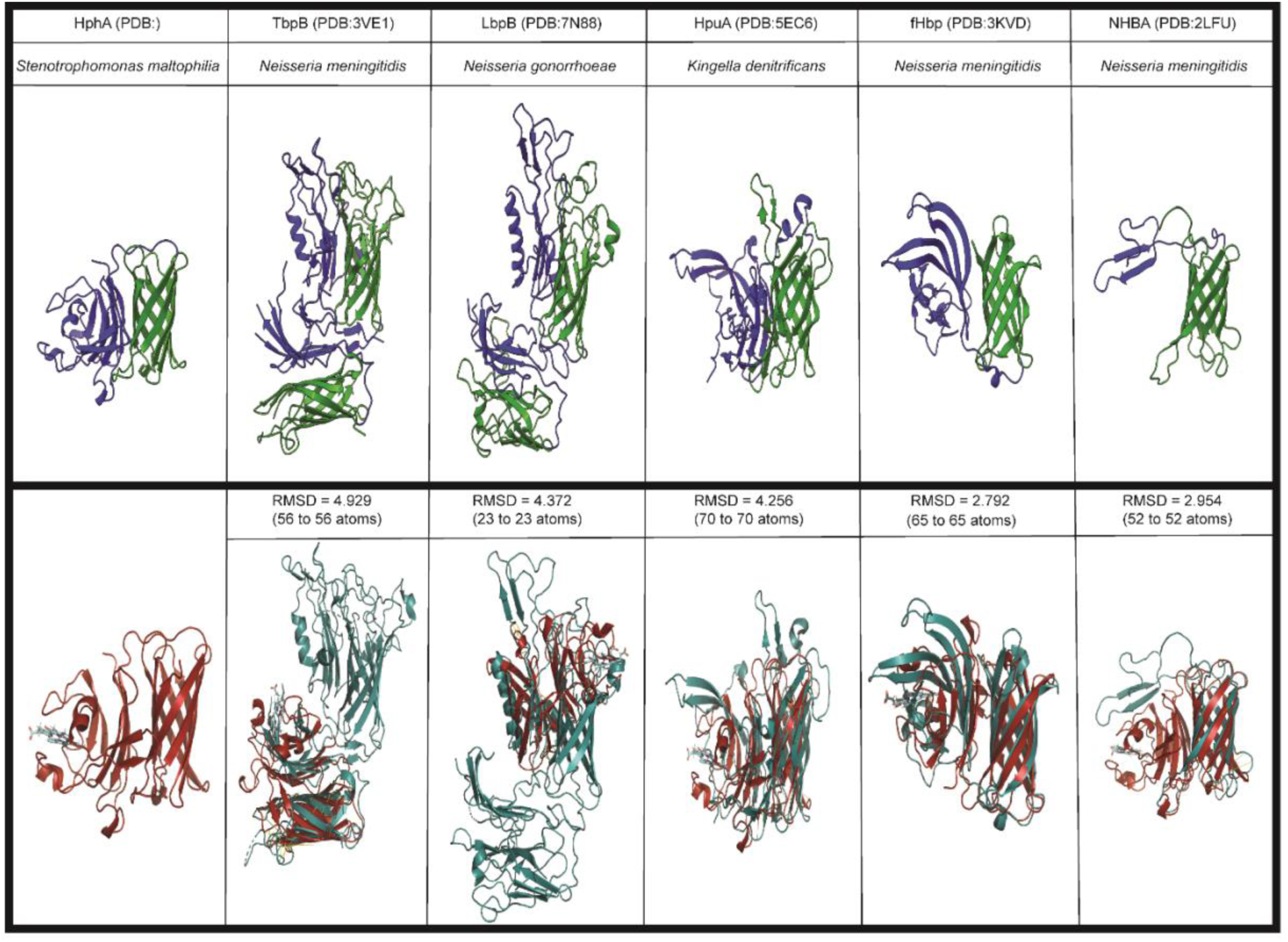
The eight stranded anti-parallel β-barrel domain (green) of HphA is structurally similar to surface lipoproteins in Neisseria spp.. TbpB (PDB: 3VE1) and LbpB (PDB:7N88) from *N. meningitidis* and *N. gonorrhoeae*, respectively, contains two β-barrel and β-handle (blue) domains. As of now, only the full length structure of HpuA (PDB:5EC6) from K. denitrificans is solved. However, the structure of HpuA in *N. gonorrhoeae* is available with only its C-terminal β-barrel (PDB:5EE2). HpuA, fHbp (PDB:3KVD), and NHBA (PDB:2LFU) contain single copies of β-barrel. Figures in the top panel were generated using ChimeraX^57^. Structural alignment was performed between the *S. maltophilia* HphA and five surface lipoproteins. The β-barrel, corresponding to residue #130-232, of *S. maltophilia* HphA was aligned with β-barrel of each surface lipoproteins. The root-mean-square deviation (RMSD) was generated using PyMOL^58^.

**Supplementary Figure 2.**
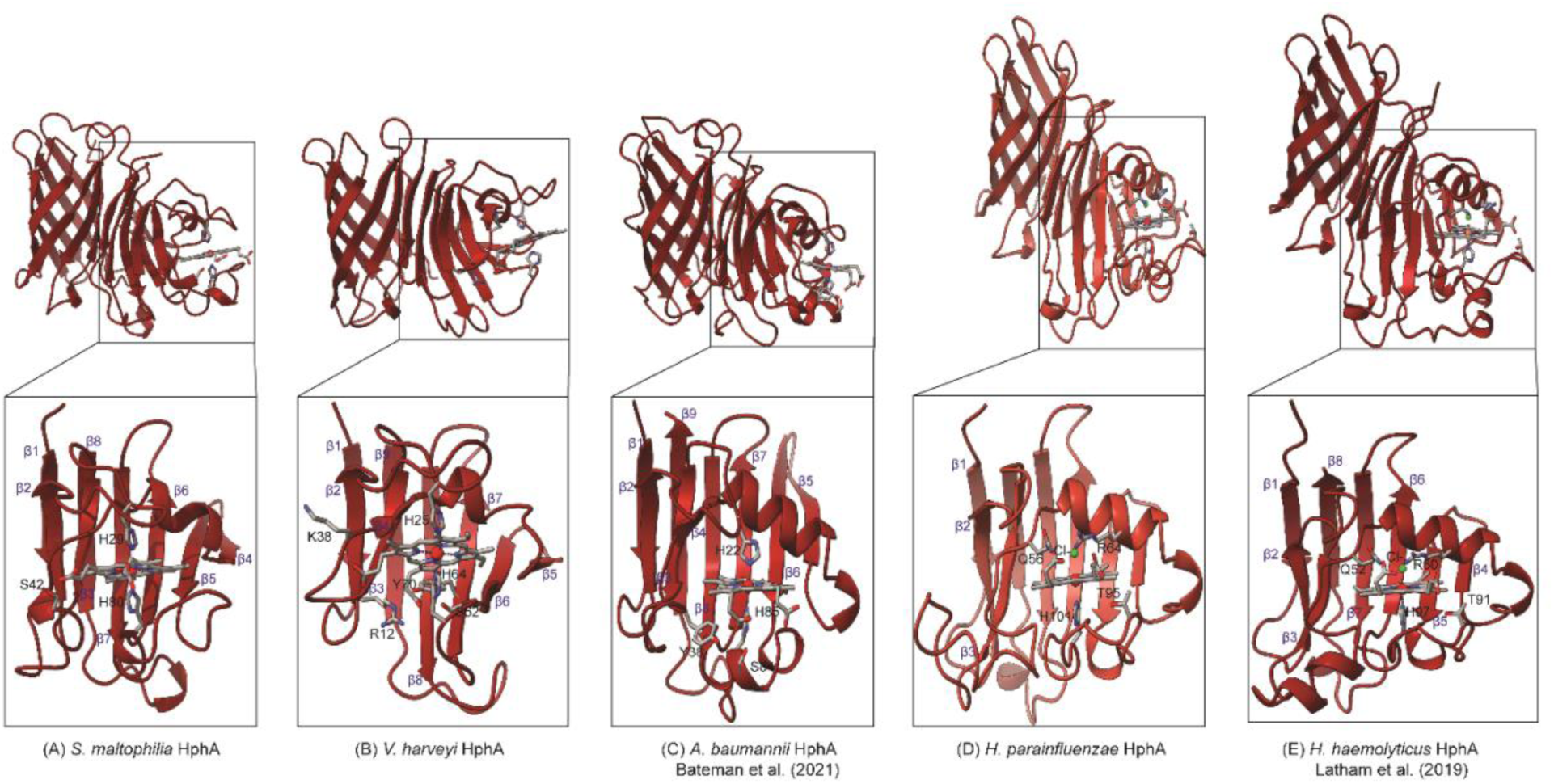
Five structurally determined HphA from different bacterial species: (A*) S. maltophilia*, (B) *V. harveyi*, (C) *A. baumannii*, (D) *H. parainfluenzae*, (E) *H. haemolyticus*. The N-terminal heme binding region, highlighted in the box, is rotated 60° and zoomed in based on the y-axis. Residues interacting with the heme ligand are indicated with the single amino acid residue code and number. The number of the amino acid residues are based on the mature protein sequences. Chloride ion and ferric iron is indicated in lime green and red, respectively.

**Supplementary Figure 3.**
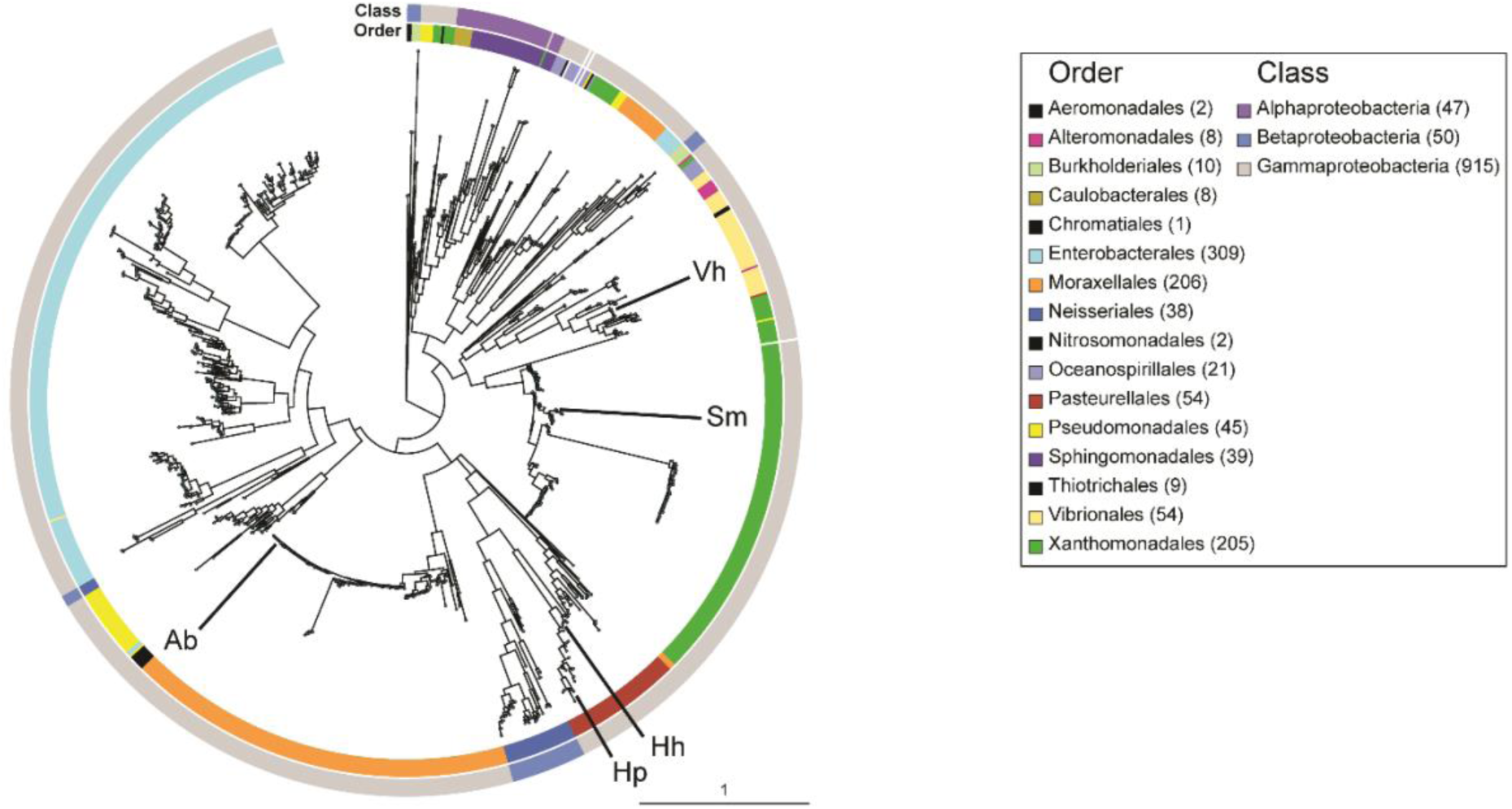
Homologs of HphAs in Proteobacteria from the ‘strict set’. The strict set contains a total of 1212 unique sequences. The inner banner indicates the order while the outer banner surrounding the tree represents the class. Five structurally identified HphAs are indicated in the arrows. The annotation Ab, Hp, Hh, Vh, Sm indicates *A. baumannii*, *H. parainfluenzae*, *H. haemolyticus*, *V. harveyi*, and *S. maltophilia*, respectively. The numbers inside the brackets indicates the number of unique accessions found in each class or order. The trees were generated using RAxML.

**Supplementary Figure 4.**
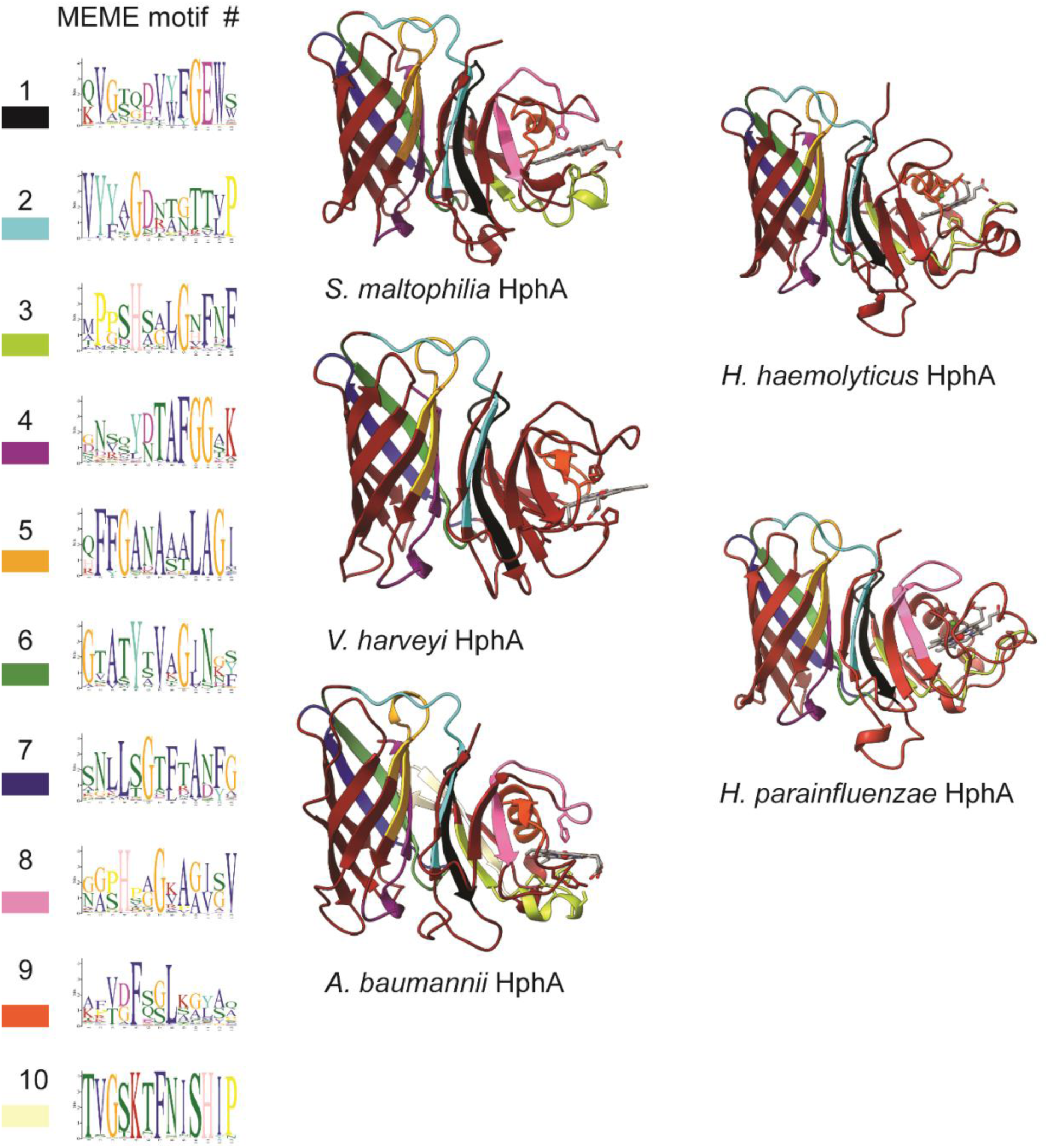
MEME analysis of HphAs. The mature protein sequence of 540 HphA homologs were analyzed to identify shared motifs. The most prevalent Motif-1 was found in β7 strand in *S. maltophilia*, *H. parainfluenzae*, and *H. haemolyticus* and β8 strand in *V. harveyi* and *A. baumannii*. The heme coordinating residues were mainly found in motif #3 with an exception of *V. harveyi* which did not contain Motif-3 but instead, contain Motif-9. The presence of Motif-1 and Motif-3/Motif-9 was used as a filtering step to generate the ‘strict set’ of HphA homologs.

**Supplementary Figure 5.**
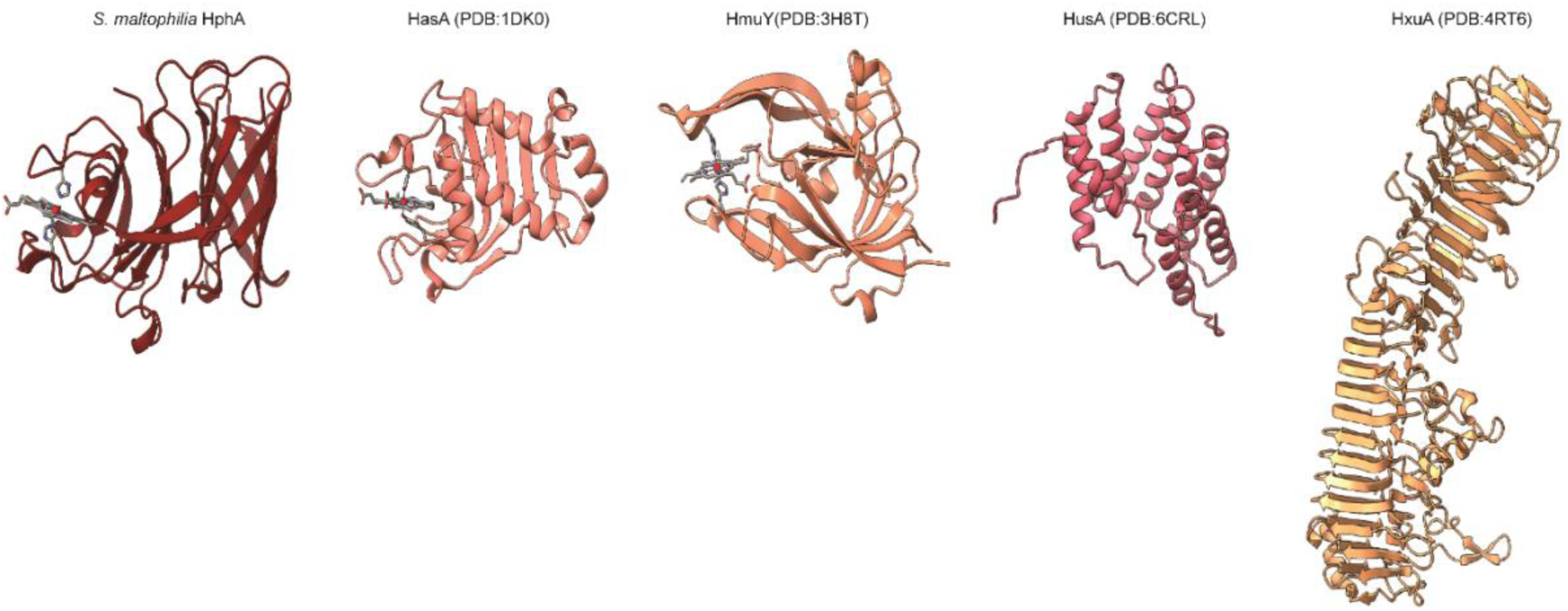
Known hemophore structures of Gram-negative bacteria. HphAs have a different overall fold compared to existing Gram-negative hemophores. HasA contains a mixed α/β fold resembling a fish capturing the heme. HmuY mimics a right hand with its β-strands and traps the heme between what resembles the thumb and fingers. HusA forms a right-handed super-helical structure with its α-helices and binds heme in its hydrophobic groove. HxuA is the largest type of an unconventional hemophore that cannot bind heme directly but scavenges heme from hemopexin. HxuA can either be anchored or secreted. HphA is the largest hemophore among the conventional hemophores.

**Supplementary Figure 6.**
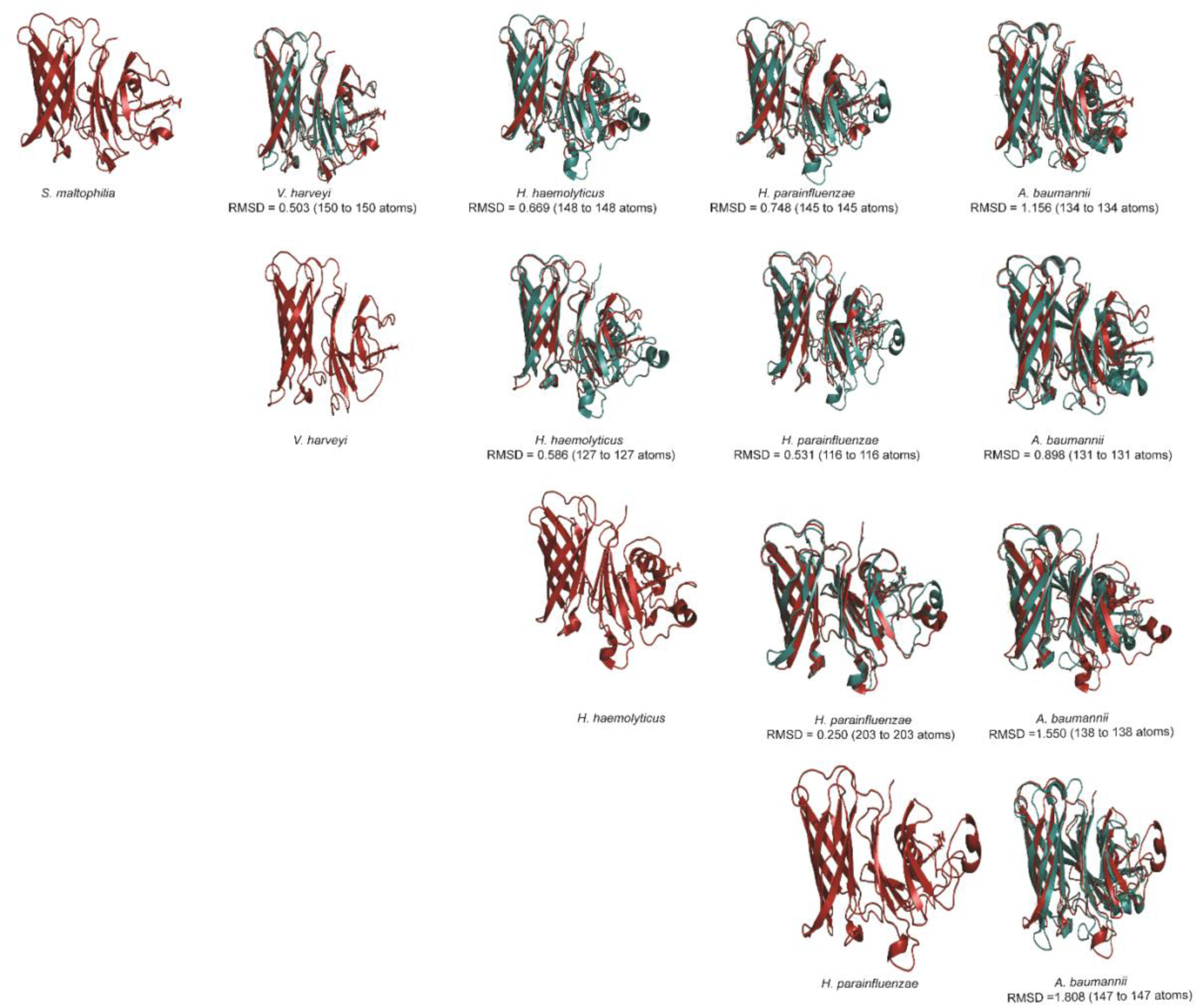
Structural similarities between the HphAs solved to date. The residues coloured in teal were aligned with the reference HphA coloured in firebrick red. Root-mean-square deviation (RMSD) was calculated via PyMOL^58^.

**Table 1.** Sequence identity and similarity of HphAs. The sequence identity and similarity of each HphAs were calculated using ‘Ident and Sim’ program of ‘The Sequence Manipulation Suite’ (www.bioinformatics.org/sms/)^59^.

*A. baumannii* HphA vs Other HphAs
|  | *S. maltophilia* | *V. harveyi* | *H. haemolyticus* | *H. parainfluenzae* |
| --- | --- | --- | --- | --- |
| Alignment length | 281 | 280 | 295 | 298 |
| Identical residues | 110 | 95 | 84 | 84 |
| Similar residues | 40 | 37 | 40 | 41 |
| Percent identity | 39.15 | 33.93 | 28.47 | 28.19 |
| Percent similarity | 53.38 | 47.14 | 42.03 | 41.95 |

*S. maltophilia* HphA vs Other HphAs
|  | *V. harveyi* | *H. haemolyticus* | *H. parainfluenzae* |
| --- | --- | --- | --- |
| Alignment length | 269 | 290 | 292 |
| Identical residues | 114 | 94 | 100 |
| Similar residues | 30 | 50 | 40 |
| Percent identity | 42.38 | 32.41 | 34.25 |
| Percent similarity | 53.53 | 49.66 | 47.95 |

*V. harveyi* HphA vs Other HphAs
|  | *H. haemolyticus* | *H. parainfluenzae* |
| --- | --- | --- |
| Alignment length | 284 | 288 |
| Identical residues | 80 | 87 |
| Similar residues | 45 | 34 |
| Percent identity | 28.17 | 30.21 |
| Percent similarity | 44.01 | 42.01 |

*H. haemolyticus* HphA vs Other HphAs
|  | *H. parainfluenzae* |
| --- | --- |
| Alignment length | 277 |
| Identical residues | 183 |
| Similar residues | 33 |
| Percent identity | 66.06 |
| Percent similarity | 77.98 |

**Table 2.** Data collection and refinement statistics of S. maltophilia HphA.

| Item | Value |
| --- | --- |
| Resolution range | 48.25 - 1.8 (1.864 – 1.8)* |
| Space group | P12<sub>1</sub>1 |
| Unit cell | a = 46.618, b= 76.663, c=62.264 |
| Multiplicity | 5.2 (2.0) |
| Completeness (%) | 82.34 (36.41) |
| Mean I/sigma(I) | 13.86 (1.57) |
| Wilson B-factor | 19.68 |
| R-merge | 0.08213 (0.3273) |
| R-meas | 0.09002 (0.4158) |
| CC1/2 | 0.997 (0.829) |
| CC* | 0.999 (0.952) |
| R-work | 0.1534 (0.2447) |
| R-free | 0.1800 (0.3145) |
| Protein residues | 464 |
| RMS (bonds) | 0.007 |
| RMS (angles) | 0.91 |
| Ramachandran favoured (%) | 98.26 |
| Rotamer outliers (%) | 0.63 |
| Clash score | 2.02 |
| Average B-factor | 24.79 |

\*Values in parentheses are for the highest-resolution shell

**Table 3.**
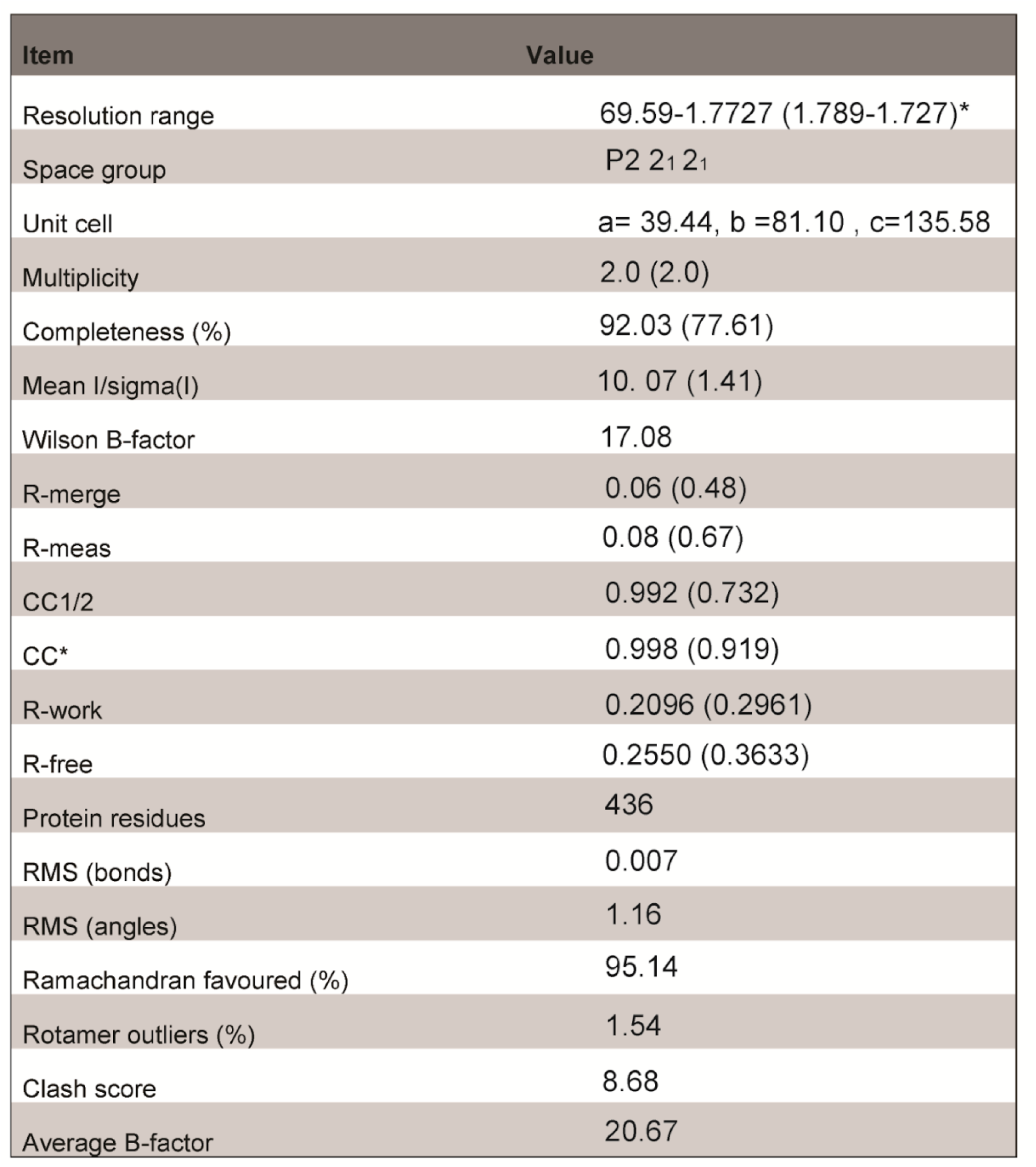
Data collection and refinement statistics of *V. harveyi* HphA

**Table 4.**
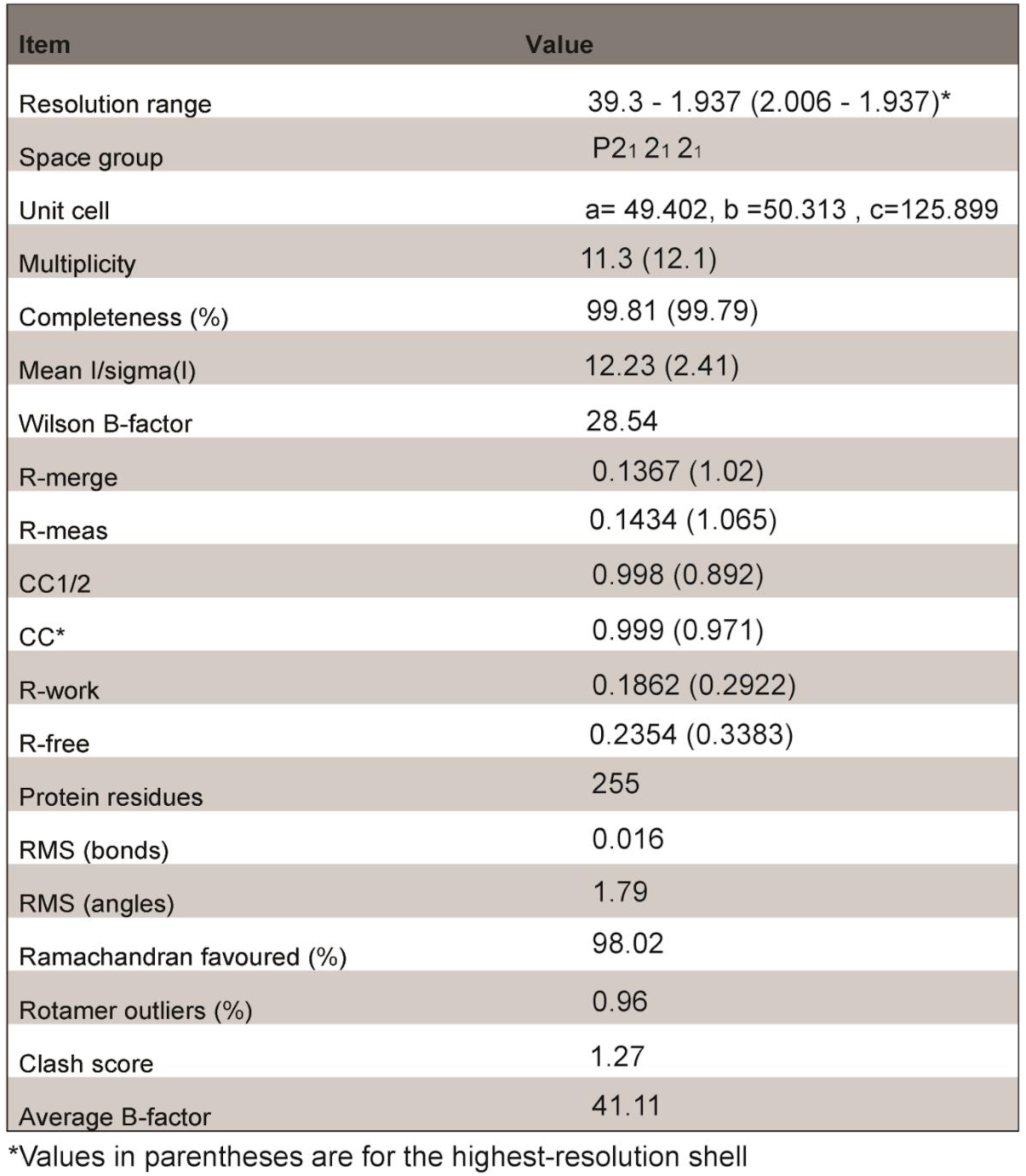
Data collection and refinement statistics of *H. parainfluenzae* HphA

| Item | Value |
| --- | --- |
| Resolution range | 39.3 - 1.937 (2.006 - 1.937)* |
| Space group | P2<sub>1</sub> 2<sub>1</sub> 2<sub>1</sub> |
| Unit cell | a= 49.402, b =50.313 , c=125.899 |
| Multiplicity | 11.3 (12.1) |
| Completeness (%) | 99.81 (99.79) |
| Mean I/sigma(I) | 12.23 (2.41) |
| Wilson B-factor | 28.54 |
| R-merge | 0.1367 (1.02) |
| R-meas | 0.1434 (1.065) |
| CC1/2 | 0.998 (0.892) |
| CC* | 0.999 (0.971) |
| R-work | 0.1862 (0.2922) |
| R-free | 0.2354 (0.3383) |
| Protein residues | 255 |
| RMS (bonds) | 0.016 |
| RMS (angles) | 1.79 |
| Ramachandran favoured (%) | 98.02 |
| Rotamer outliers (%) | 0.96 |
| Clash score | 1.27 |
| Average B-factor | 41.11 |

\*Values in parentheses are for the highest-resolution shell

